# Longitudinal impact of oral probiotic *Lactobacillus acidophilus* vaccine strains on the immune response and gut microbiome of mice

**DOI:** 10.1101/691709

**Authors:** Zaid Abdo, Jonathan LeCureux, Alora LaVoy, Bridget Eklund, Elizabeth Ryan, Gregg Dean

## Abstract

The potential role of probiotic bacteria as adjuvants in vaccine trials led to their use as nonparenteral live mucosal vaccine vectors. Yet, interactions between these vectors, the host and the microbiome are poorly understood. This study evaluates impact of three probiotic, *Lactobacillus acidophilus*, vector strains, and their interactions with the host’s immune response, on the gut microbiome. One strain expressed the membrane proximal external region from HIV-1 (MPER). The other two expressed MPER and either secreted interleukin-1β (IL-1β) or expressed the surface flagellin subunit C (FliC) as adjuvants. We also used MPER with rice bran as prebiotic supplement. We observed a strain dependent, differential effect suggesting that MPER and IL-1β induced a shift of the microbiome while FliC had minimal impact. Joint probiotic and prebiotic use resulted in a compound effect, highlighting a potential synbiotic approach to impact efficacy of vaccination. Careful consideration of constitutive adjuvants and use of prebiotics is needed depending on whether or not to target microbiome modulation to improve vaccine efficacy. No clear associations were observed between total or MPER-specific IgA and the microbiome suggesting a role for other immune mechanisms or a need to focus on IgA-bound, resident microbiota, most affected by an immune response.

## Introduction

The relationship between the microbiome and its host has been rigorously studied during the last decade resulting in evidence supporting its role in health and disease^1,2,3,4,5^. It was also established that the microbiome is dependent on diet and on its environment and could be modulated, for better or worse, by use of antibiotics, probiotics and/or prebiotics^6,7,8,9,10,11^. Use of probiotics has been shown to impact the gut microbiome due to direct competition or cooperation with the resident microbiota, and was shown to have a direct effect on the host through immune modulation and enhanced barrier function^12,13,14,15^.

It is clear that the microbiome greatly influences mucosal health^3,16^, but how vaccines, and any subsequent mucosal immune response, influence the microbiome is poorly understood^17^. There is increasing evidence that the immunogenicity and efficacy of current vaccines are related to the intestinal microbiome^18,19,20^. Multiple studies have also shown that probiotic administration prior to or concurrent with vaccination enhances B cell and antibody responses and provides the mucosa with direct protection from infection through interactions with the innate immune system^21,20,22,23^. Trials of both parenteral and nonparenteral vaccines, in conjunction with probiotic administration, also point to probiotic bacteria as adjuvants^17^. The mechanisms behind this phenomenon are incompletely understood but are likely due to probiotic surface structures^24^. This inherent adjuvanticity has led to the use of probiotics, typically lactic acid bacteria, as nonparenteral live mucosal vaccine vectors^25,26,27^.

Species and subspecies of the genus *Lactobacillus* are an important and heavily studied group of Gram-positive lactic acid bacteria used for food preservation, food bioprocessing, and as probiotics. Most lactobacilli possess acid and bile salt tolerance, allowing them to survive the hostile environment of the stomach and proximal duodenum^28,29,30^. Additionally, several cell surface components of lactobacilli are recognized by immune cells via pattern recognition receptors (PRR)^31^. In particular, lipoteichoic acid (LTA), peptidoglycan (PG), and muramyl dipeptide (the subcomponent of PG) are the major immune stimulators recognized by the heterodimeric Toll-like receptor (TLR) 2/6 and nucleotide-binding oligomerization domain 2 (NOD2), respectively^32,33,34^. This capacity to interact with the innate immune system helps explain why some species of lactobacilli are effective inducers of mucosal antibodies, especially IgA^35^. The probiotic strain *Lactobacillus acidophilus* NCFM is particularly promising as an oral vaccine vector for several reasons: (1) immune stimulation via PRRs as was just described, as well as binding to dendritic cells (DCs) through DC-specific intercellular adhesion molecule 3 (ICAM-3)-grabbing nonintegrin (DC-SIGN)^36^, (2) acid and bile tolerance^29,30^, and (3) expression of mucus-binding proteins and association with the mucosal epithelium^37,38^.

In this study we evaluated the impact of *Lactobacillus acidophilus*, as a probiotic vaccine vector, and its interactions with the host’s immune response, on the gut microbiome. We utilized three modified *L. acidophilus* strains with different constitutive adjuvants. All three strains expressed the membrane proximal external region (MPER) from Human Immunodeficiency Virus 1 (HIV-1) within the context of the major Surface-layer protein A (SlpA) that was developed in previous work^39^. The MPER epitope alone is a very weak B-cell immunogen, so to increase immunogenicity the two additional *L. acidophilus* vaccine strains were modified to either secrete soluble interleukin-1β (IL-1β, an inflammatory cytokine) or surface-expressed flagellin protein C (FliC, a TLR5 agonist). Both of these adjuvant strains were previously identified for increasing immunogenicity against MPER^39,40,41^. In addition, we used the MPER-expressing *L. acidophilus* (no IL-1β or FliC) along with rice bran as a prebiotic supplement. Rice bran has previously shown adjuvant properties to rotavirus vaccination in pigs and to enhance growth of probiotics^42^. To our knowledge no probiotic vaccine has been tested for gut microbiome alterations, and prior evidence with other oral vectors suggests that oral vaccines do not cause significant perturbations to the host microbiome, unlike what we have observed.

## Results

### Vaccination-Induced Differences in Alpha Diversity

Results of model fitting highlighted differences in Chao1 and Shannon diversity measures between the vaccine treatments over time (Figure 1 and Table S1). Vaccine treatments used included *Lactobacillus acidophilus* (LA) expressing MPER as vaccine vector (MPER), LA expressing MPER and secreting interleukin-1β as adjuvant (IL1b), LA expressing MPER and also expressing surface flagellin subunit C as adjuvant (FliC), LA expressing MPER and a custom rice bran diet as prebiotic (RB), LA wild-type strain as positive control (WT), and a negative control group (NG) (*Materials and Methods*). The model describing Chao1 richness (Table S1) highlights variation between treatments and between mice within the same treatment—it also includes a nonlinear temporal trend driven by the FliC and RB treatments (Figure 1A). FliC witnessed an increase in richness for weeks 2 through 6 that was mostly similar to that of NG and WT but significantly different from that of IL1b and RB for all three time points and from that of MPER for time points 2 and 4. FliC richness declined during weeks 8 and 10 and was similar to that observed for RB for week 10, which was qualitatively lower than all other treatments. Richness of the RB treatment was lower than all other treatments starting at week 2 and continuing to the end of the experiment. In addition to being significantly lower than that of FliC, RB richness was also significantly lower than that of NG and WT during weeks 2 through 6 highlighting the combined impact of a change in the microbial environment induced by both vaccine application and the presence of the rice bran prebiotic in the modified diet. The upward trend after the 4^th^ week could be interpreted as a possible recovery in richness as the mice adjusted to the change in diet and imposition of the vaccine. In general, IL1b and MPER had a similar trend and so did NG, FliC and WT, with NG having the lowest slope of increase in richness over time. IL1b had the second lowest Chao1 richness besides RB, also indicating a putative compound effect of an immune response due to the expression of IL-1β and the impact of the vaccine vector. No significant differences were observed between the treatment levels at time 0 (highlighted by overlap between the 95% credibility intervals at that time point). Qualitatively, though, the negative control (NG) treatment showed somewhat higher richness than all other treatments followed by both FliC and the positive control (WT) at that point in time. All treatments seemed to converge to the same richness level by the end of the experiment.

**Figure 1:**
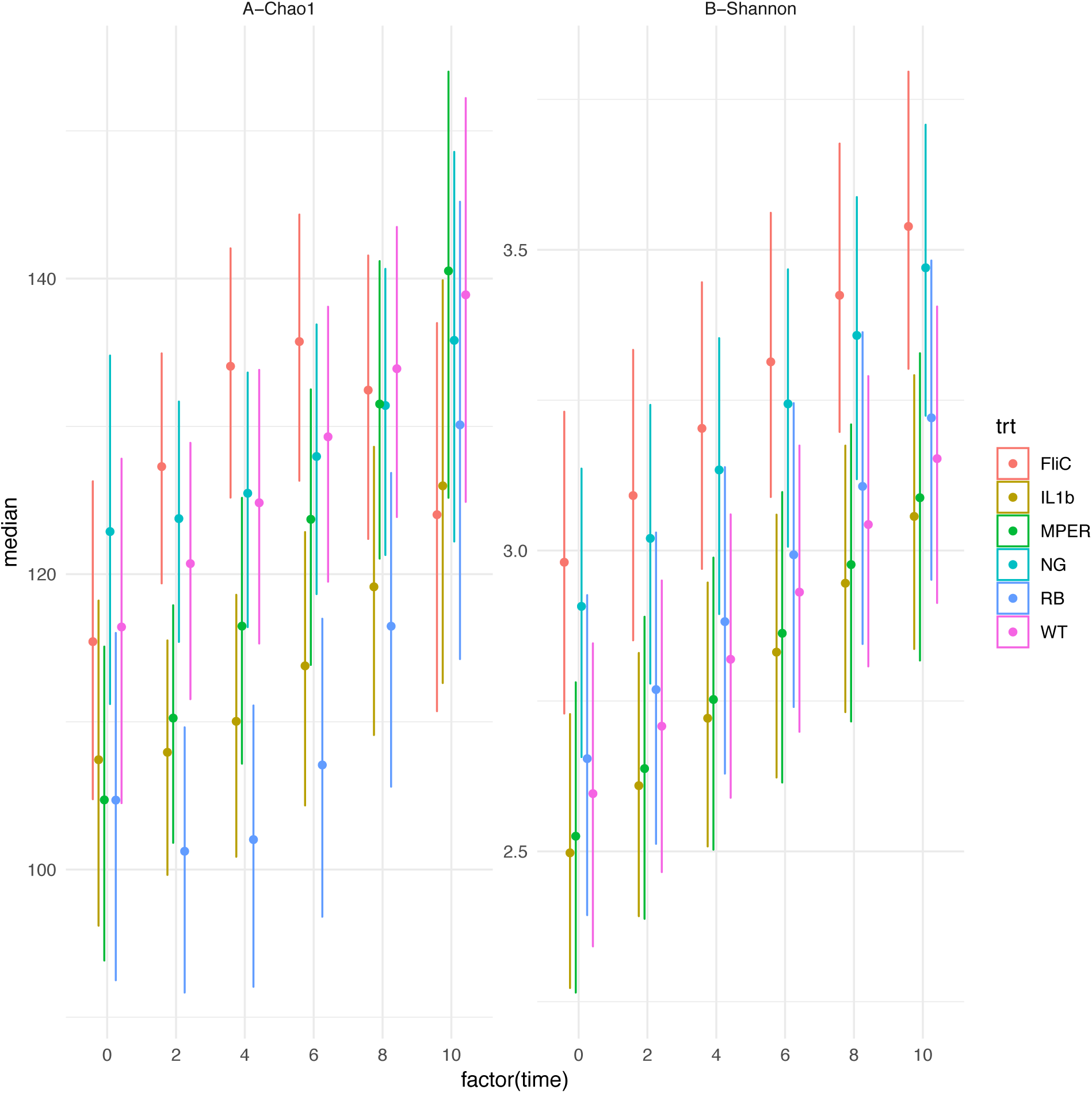
Change of alpha diversity over time in association with the different treatment levels: (A) The 95% credibility intervals of the expected Chao1 richness per time point (0, 2, 4, 6, 8 and 10) under each of the treatment levels. (B) The 95% credibility intervals of the expected Shannon diversity index per time point (0, 2, 4, 6, 8 and 10) under each of the treatment levels. Both panels are based on the observed fecal samples.

Shannon diversity index accounts for both richness and abundance of the different OTUs within the observed samples. A simple linear regression model, with intercepts that varied per-treatment and per-mouse, was selected to describe the trends in this index (Figure 1B and Table S1). Similar to Chao1, this model highlights differences due to the treatments and variation between mice within each treatment. Figure 1B highlights an increased diversity of the FliC treatment, compared to all treatments other than NG, that was significantly different from that under the MPER and the IL1b treatments, similar to what has been observed for Chao1. Shannon’s diversity for FliC seems to be similar to that of the negative control group. Both the MPER and the IL1b treatments were very similar and have the lowest diversity compared to all other treatment levels though not significantly different from the WT and RB treatments. Figure 1B indicates a linear increase in diversity as time progresses in the experiment. This increase might be a result of the mouse acclimation to the stress imposed by the gavage process used for vaccination.

Figure S2 compares the alpha diversity indexes between the different treatment levels for the cecal samples. The model identified to best fit these data did not show significant differences between the observed microbiomes, reflected in the overlap between the 95% credibility intervals observed in the figure. However, it is worth noting that, qualitatively, some of the trends observed in the fecal samples were also found in the cecal samples including the increased richness and diversity under the FliC treatment and the reduced richness under the RB treatment.

### Vaccination-Induced Trends in Beta Diversity

Separation between microbiomes associated with each treatment level is clear in the NMDS presented in Figure 2. An exception was MPER and IL1b, which seemed to overlap, corroborating the trends observed in the alpha diversity analyses above. Maximum separation was observed between the microbiome associated with the RB treatment indicating a possible compound effect of the prebiotic on the response of the microbiome to vaccination, also similar to above. Similar trends were observed for the cecal samples (Figure S3) where MPER and IL1b, and the NG, seemed to be closer together than those of the other treatment levels.

**Figure 2:**
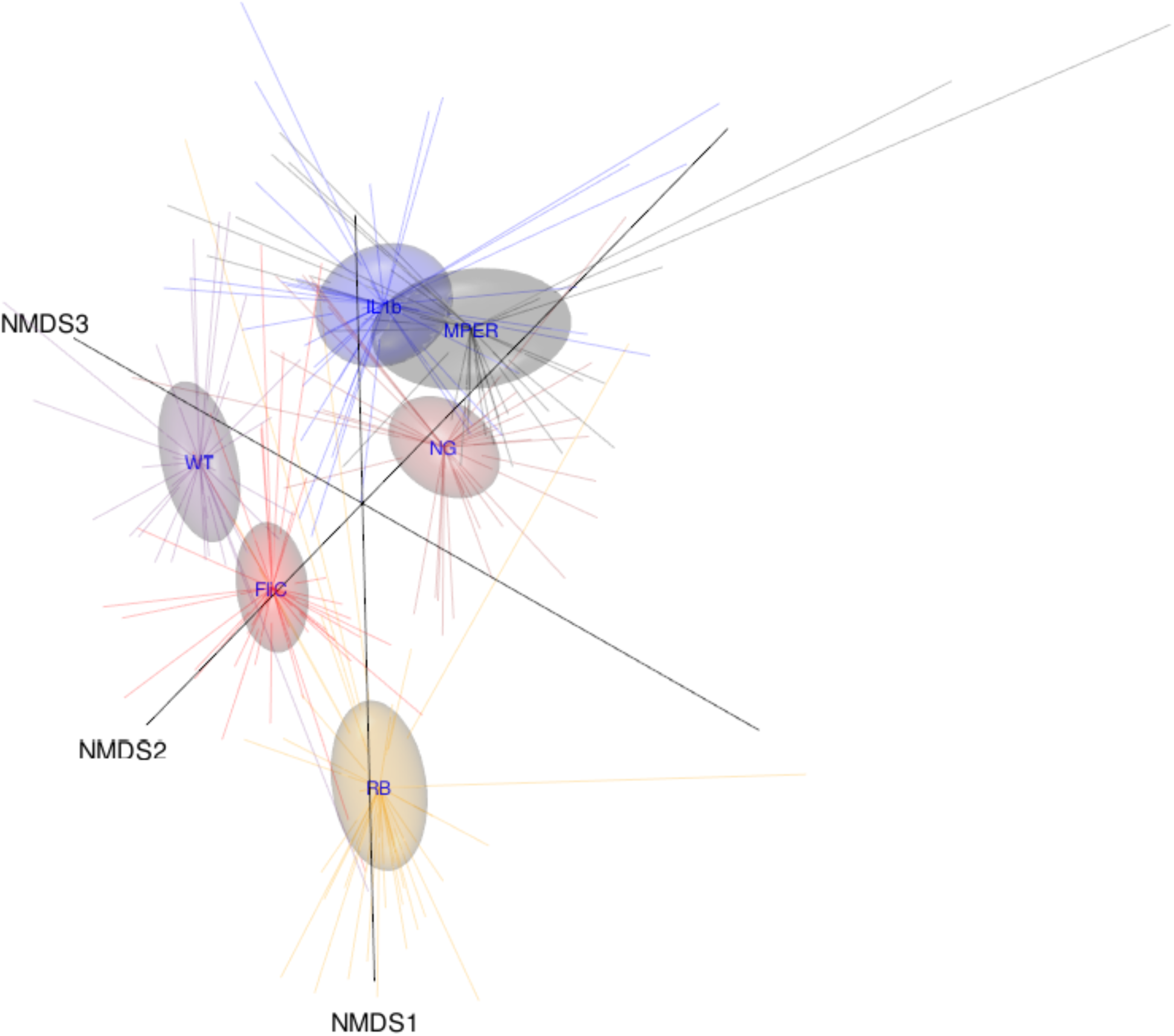
**Three-dimensional (3D) nonmetric multidimensional scaling (NMDS) ordination plot** of the beta diversity of fecal samples measured using the Bray-Curtis distance and aggregated by treatment level. The figure highlights the different data points as tips of the star shapes emitted from the centroids representing the treatment levels and the associated 95% confidence ellipsoids.

Figure 3 provides a two-dimensional view of the trends of the beta diversity per-treatment. The figure indicates shifts in the microbial community structure occurring over time. This shift is clearly observed for the WT, MPER, and IL1b treatments where the microbial community seems to progress from its state at week 0 to a different state at week 10. This shift occurs in one step for the MPER treatment, where weeks 4-10 cluster at a new equilibrium, but gradually for the WT and the IL1b treatments. Although we observed separation between week 0 and week 10 for the FliC treatment, all time points clustered toward time zero. This observation is in concordance with the alpha diversity, indicating that vaccination supplemented with FliC has the least impact on the microbial community of the host. The RB treatment showed a profound jump in the microbial community structure from its state at time −1, one week prior to application of the treatment and the rice bran diet, and then to a new equilibrium state after progressing through the first treatment application at week 0. The negative control group did not show any significant trends as expected.

**Figure 3:**
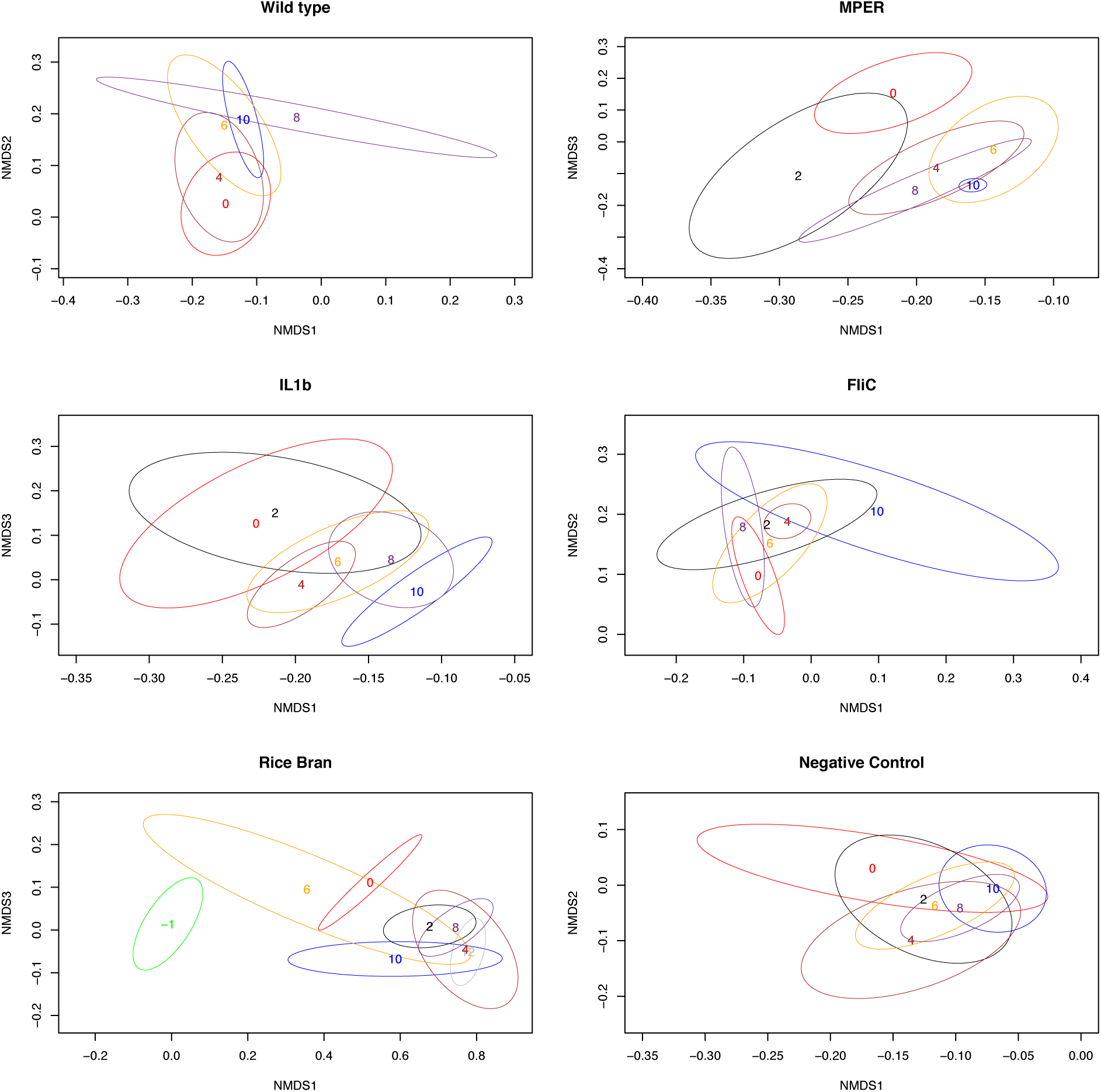
**Per-treatment NMDS ordination plots** representing beta diversity changes over time per treatment level. Each number marks the centroid of the 95% confidence ellipsoid at the time point the number represents (0, 2, 4, 6, 8 or 10).

### Taxonomic Level Trends and Association with treatment groups

There were ten phyla observed within the microbial community structure within all mice under the different treatments for all time points. Five of these phyla were found to have relative abundance larger than 1% per-sample and these are: Firmicutes, Bacteroidetes, Tenericutes, Actinobacteria and Proteobacteria—presented in Figure 4 for the fecal samples and in Figure S4 for the cecal samples. Actinobacteria was only detected in the negative control while Proteobacteria was observed in the MPER treatment. As expected, Firmicutes and Bacteroidetes dominated the mouse gut microbiome in all samples conforming to previous observations^4^.

**Figure 4:**
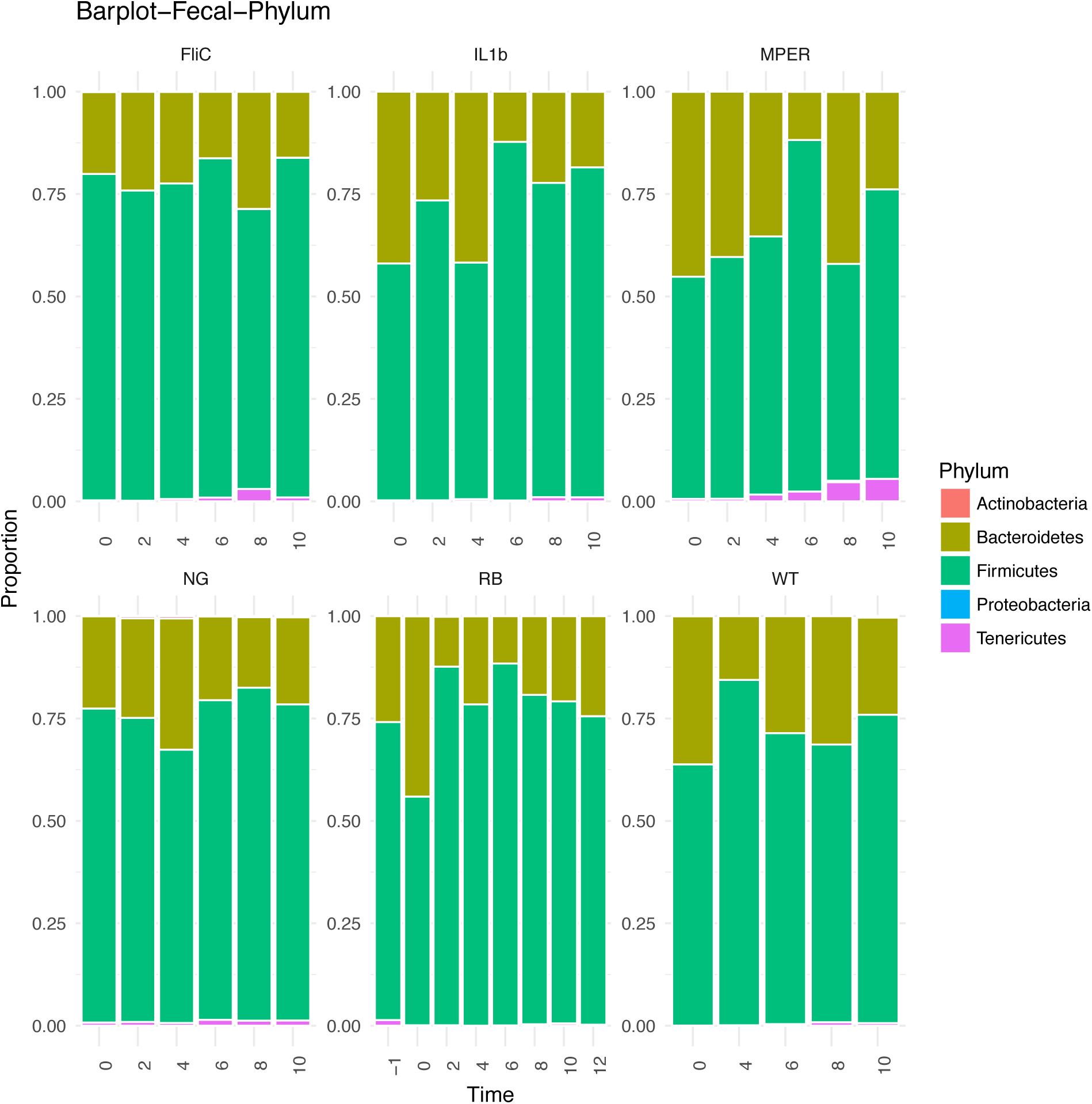
**Bar plots** representing the phylum level taxonomic distribution per treatment level per time point of the fecal samples.

Using Random Forests (RF) our goal was to identify drivers of the putative differences in response to the treatments. We added time as a feature to account for the possible temporal effect in discriminating between treatments. The optimal number of features (OTUs and time) used in constructing the decision trees in the random forest was identified to be 53 for the fecal samples and 9 for the cecal samples. This number was observed to minimize the out-of-bag (OOB) error rate (Figure S5 and *Materials and Methods*).

For the fecal samples the overall OOB estimate of error rate was found to be about 20%. That is, 80% of the time samples were assigned to their correct treatment. The OOB error rate ranged from 14.3% to 24.1% per treatment, as shown in the confusion matrix presented in Table S2, with assignment to FliC having the lowest error rate and assignment to WT having the highest. The table indicates closeness between MPER and IL1b; 6 out of 35 samples belonging to IL1b were misclassified as MPER and 6 out of 32 MPER samples were misclassified as IL1b corroborating some of the results observed in association with the alpha and beta diversity above. The table also highlights non-reciprocal association between WT and IL1b where 5 out of 29 WT samples were misclassified as IL1b with only one IL1b sample misclassified as WT. Table S3 indicates that much of the misclassification occurred at time zero, the time when the microbial richness was observed to be similar as described above.

Figure 5 shows the relative Mean Decreasing Gini importance plot of the 30 most impactful features, all of which are OTUs. Table S4 associates these OTUs with their taxonomic assignment; the higher the Gini value the more impactful is the feature. It is interesting that time was not identified as highly important for classifying the observed samples. It is also worth noting that an OTU part of the genus Bifidobacterium (phylum Actinobacter) was the most impactful discriminator. Other observed OTUs belonged to the families Lachnospiraceae (16), Ruminococcaceae (5), Clostridales_vadinBB60_group (6), Streptococcaceae (1) and Bacteroidales_S24-7_group (1) (Table S4). Ten of the Lachnospiraceae OTUs were not classified; the remaining six belonged to the genera Tyzzerella, Acetatifactor, Lachnospiraceae_UCG-001, Marvinbryantia, Lachnoclostridium and Lachnospiraceae_NK4A136_group. OTUs identified part of the Ruminococcaceae family belonged to the Ruminiclostridium, Anaerotruncus, Ruminococcaceae_UCG-014 and Oscillibacter genera. All of the taxa belonging to Clostridiales_vadinBB60_group belonged to the Clostridiales_vadinBB60_ge genus. All of these three families were part of the Firmicutes phylum. Streptococcaceae, phylum Firmicutes as well, was the only Bacilli; Lactococcus was the genus associated with the OTU observed within this family and was ranked sixth in its impact on the correct classification. The observed OTU within family Bacteroidales_S24-7_group belonged to the genus Bacteroidales_S24-7_group_ge. This was the only OTU part of the phylum Bacteroidetes that was identified to impact the correct classification.

**Figure 5:**
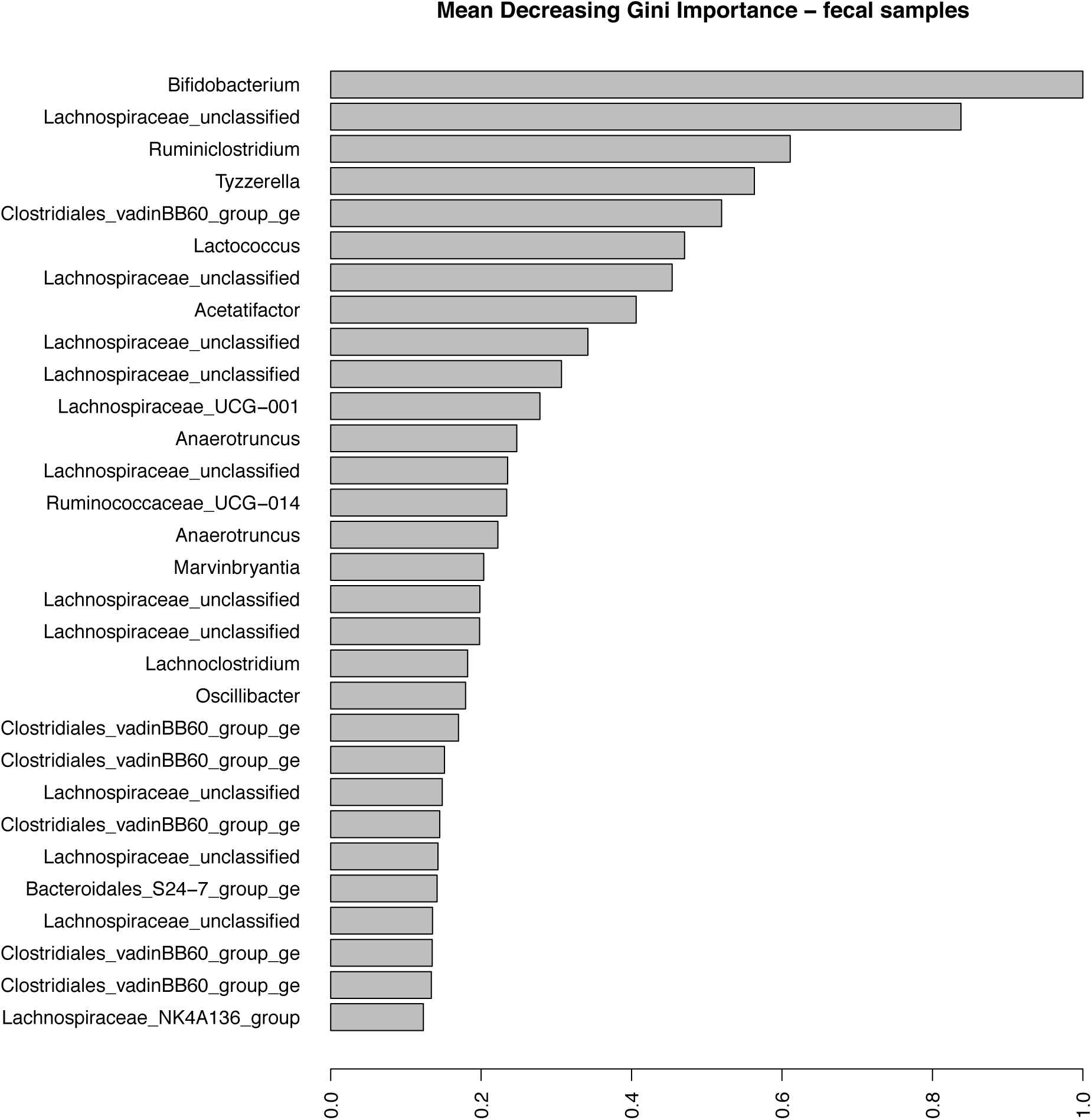
**The Mean Decreasing Gini OTU importance plot** for the fecal samples. X-axis represents the Gini importance measure where high values represent high impact of the OTUs presented on the y-axis.

Figure 6 shows the trends observed for the five most impactful taxa identified within the fecal samples. Bifidobacterium (OTU0088) was mostly observed in the NG treatment and at week 10 of the WT treatment. Lachnospiraceae_unclassified (OTU0059) was observed for the most part in association with FliC and WT. Ruminiclostridium (OTU0149), on the other hand, was mostly present in NG, MPER, IL1b and WT but not FliC or RB. Clostridiales_vadinBB60_group_ge (OTU0180) was present mostly in MPER and FliC and to a lesser degree in IL1b and NG but not in RB and WT. Tyzzerella (OTU0097) was present for the most part in RB times 0 – 12 but to a much lower degree in RB time 0 and in the other treatments. Figures describing the other 25 taxa can be found in Figure S6.

**Figure 6:**
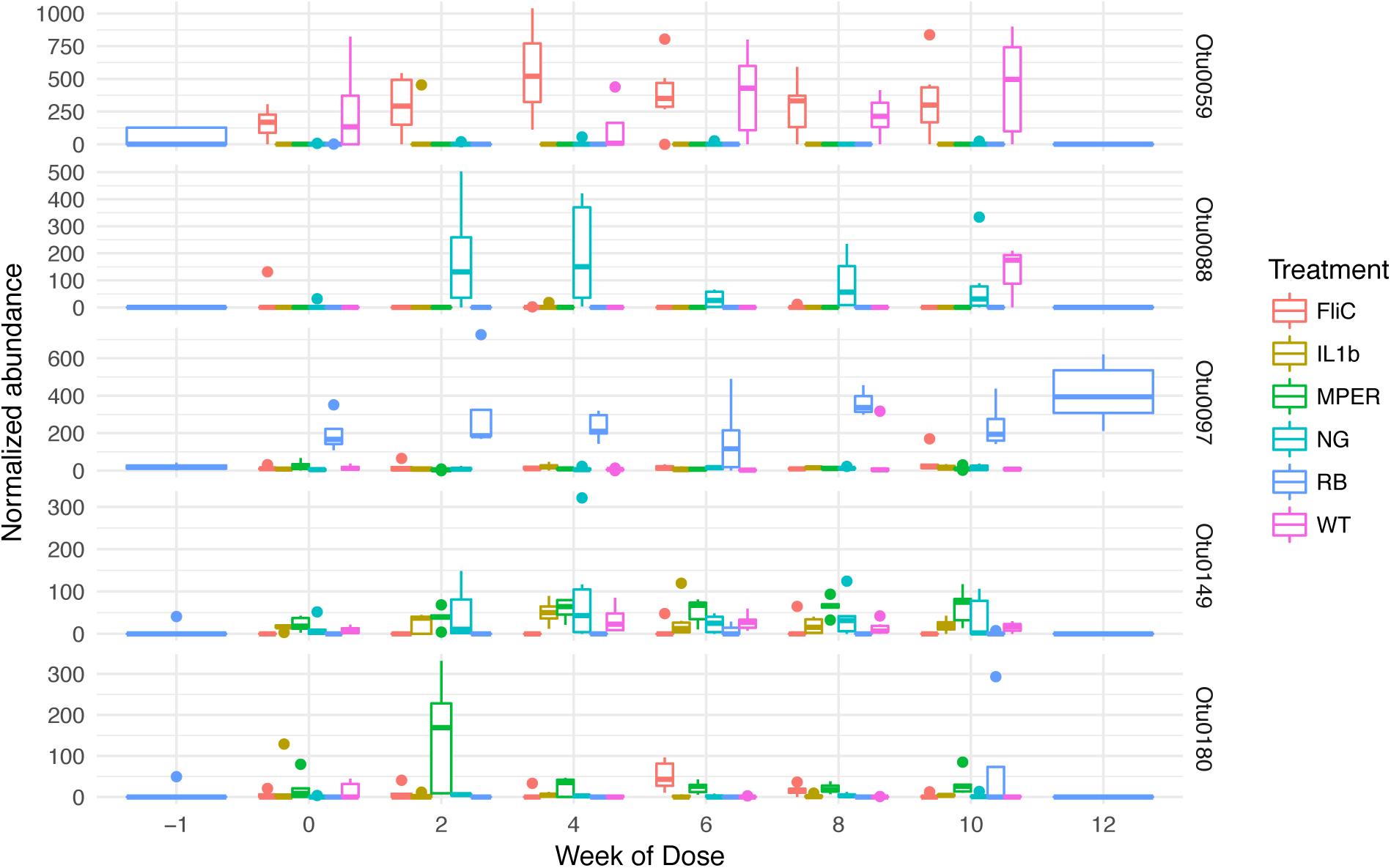
**The normalized abundance of the five most impactful OTUs** associated with the fecal samples as observed over time: Bifidobacterium (OTU0088), Lachnospiraceae_unclassified (OTU0059), Ruminiclostridium (OTU0149), Clostridiales_vadinBB60_group_ge (OTU0180) and Tyzzerella (OTU0097).

The overall OOB estimated error rate for the cecal samples was about 20.59%. That meant that samples were correctly classified to the respective treatment about 79% of the time, similar to the fecal samples. The per-treatment OOB error rate ranged from 0% for RB to 50% MPER (Table S5). This result highlights separation of the RB treatment from all others indicating possible interaction or direct impact of the rice bran components on the microbial community response to vaccination. MPER samples did not have sufficient signal to discriminate them from other samples. The OOB for all other treatments was 16.7% (1 in 6 erroneous misclassifications). Note that the small sample sizes were not conducive for accurate classification for this experiment. Figure S7 shows the relative Mean Decreasing Gini importance plot of the 30 most impactful features for the cecal samples. Table S6 associates these OTUs with their taxonomic assignments. Unlike the observed fecal samples, Bifidobacterium was not a driver of discrimination for the cecal samples. Family Lachnospiraceae was represented by 22 OTUs, 16 of which were either unclassified or uncultured; the remaining six belonged to Lachnospiraceae_NK4A136_group, Lachnospiraceae_ge, Lachnospiraceae_FCS020_group, Roseburia, Acetatifactor and Lachnoclostridium. Six taxa represented the family Ruminococcaceae and these belonged to genera Oscillibacter (2), Ruminiclostridium_6, Anaerotruncus, Ruminiclostridium_9 and Ruminiclostridium. Only one taxon belonged to the genus Lactobacillus and another to Clostridiales_vadinBB60_group_ge. The phylum Firmicutes was the only phylum observed within the top 30 impactful taxa. However, due to the near-uniformity of the Gini measure we can conclude that many of the observed taxa, and not only the top 30, had an impact on the correct classification. Many of the top 30 taxa in the cecal samples matched those highlighted in the fecal samples above.

Figure S8 shows the trends of the five most impactful taxa in the cecal samples. The top three of these OTUs (OTU0076, OTU0094 and OTU0039) were unclassified or uncultured Lachnospiraceae. OTU0076 was highly abundant in the WT samples compared to all others. OTU0094 and OTU0039 were highly abundant in IL1b, MPER and the NG samples. OTU0094 also had detectable presence in the RB community albeit at a low relative abundance. The fourth impactful OTU (OTU0012) belonged to the Lachnospiraceae_NK4A136_group genus. This genus was highly represented in the RB samples compared to all other samples. The fifth OTU (OTU0167) belonged to the Oscillibacter genus. This OTU was present in all samples with elevated relative abundance in both the IL1b and the WT treatments. Trends for the remaining 25 taxa can be found in Figure S9.

### Vaccination-Induced Trends in Total and MPER-Specific IgA

The model best describing the trend in total IgA production was one that included a treatment- and mouse-specific cubic trend and intercept, also highlighting the variability between and within treatments (Figure 7). The cubic trend was mostly driven by the FliC treatment with fluctuation of the total fecal IgA over time from high to low to high again. Worth noting is the behavior of the WT treatment where the total IgA peaked during the 6^th^ week and was significantly different than that observed during week 0 within that treatment. Total IgA production of the WT was lower than that of all other treatments at week 0. No significant correlations were observed between the total IgA and the alpha diversity measures described above (Figure S10) indicating that effectors other than total IgA might be responsible for the changes observed in the diversity of the microbiome.

**Figure 7:**
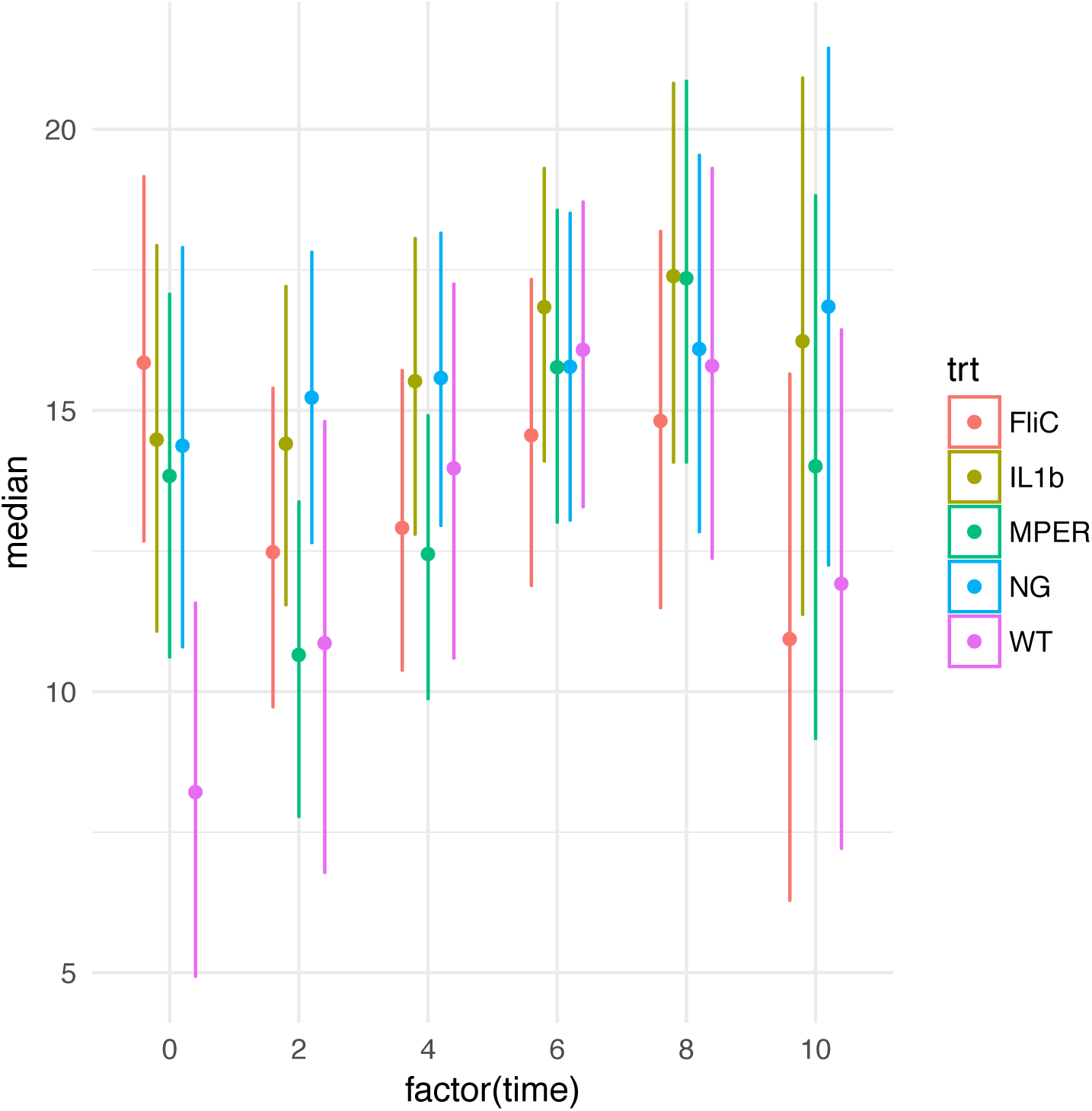
**The 95% credibility intervals of the expected total IgA per time point** (0, 2, 4, 6, 8 and 10) under each treatment level. Center points represent the median total IgA per time point.

Figure S11 shows an increase in MPER-specific IgA over time in MPER, FliC and IL1b, all of which include the MPER epitope. This is in accordance with our expectations. The figure indicates that the most pronounced induction of this response was observed in treatment IL1b. Still, some of the mice in these experiments didn’t have a detectable level of this specific IgA. No significant correlations were observed between these trends and measures of microbial diversity (Figure S10). A lack of MPER-specific IgA in WT and NG groups shows induction of this antibody was due to our vaccine and not non-specific antibody production.

### Minimal Associations Between Microbial Drivers and IgA

There were no IgA measurements observed for the cecal samples or the RB treatment. Samples with missing IgA measurements were dropped. The presented analysis is for total IgA and antigen-specific IgA samples separately, to minimize the missing data removal. Bars in color represent significance at the 0.1 cutoff p-value, without correction for multiple testing, of positive (red) and negative (blue) Spearman’s correlation between normalized taxonomic abundance of the 30 most impactful taxa and total IgA (Figure 8 and Table S7) and MPER-specific IgA (Figure S12 and Table S8). There were no particular patterns observed in these correlations between OTUs and total IgA save possibly one with Clostridiales_vadinBB60_group_ge. Clostridiales_vadinBB60_group_ge seemed to be positively correlated with IgA in all types of vaccine applications, significantly at the FliC treatment level. On the other hand, it seemed to either be neutral (WT treatment) or negatively associated with total IgA (NG treatment). Mixed results were also observed for antigen-specific IgA with most of the high correlation observed between taxa in the MPER treatment. These results are not conclusive though might support our observation that factors other than IgA might be at play in driving the microbiome’s response to vaccination or that a more targeted approach, focusing more on the resident microbiota that are most affected by the immune response, is required to better assess these associations.

**Figure 8:**
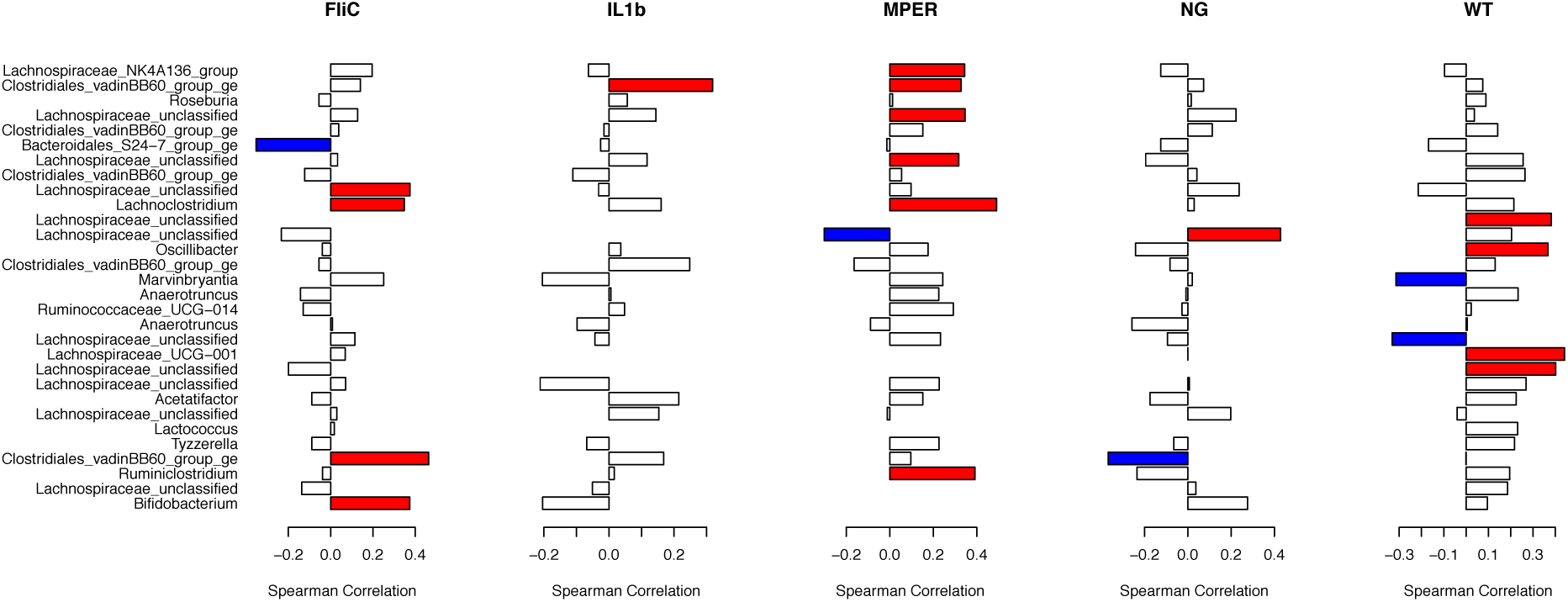
Spearman correlation plots linking the 30 most impactful taxa observed with the fecal samples and total IgA per treatment. Red and blue represent significant positive and negative correlations, respectively, at the 0.1 level of significance with no correction for multiple testing.

## Discussion

In this paper we studied the impact of an oral vaccine vector on the murine gut microbiome, utilizing the probiotic *L. acidophilus* induced immune response. We evaluated this impact using an HIV-specific peptide vaccine (MPER) augmented with either FliC or IL-1β as adjuvants and compared its use against a regimen that included rice bran as a prebiotic, along with positive and negative controls.

Maintenance of alpha and beta diversity was observed in association with the FliC treatment as compared to the negative control. This treatment utilized recombinant *L. acidophilus* (rLA) allowing for co-expression of the MPER epitope and the *Salmonella enterica* serovar Typhimurium flagellin (FliC). Maintenance of diversity could result when there is no effect of rLA on the microbiome directly through competition and/or indirectly through immune response. Our previous work has shown that FliC is an effective adjuvant that improves the response to the vaccine by stimulating activity of the Toll-like receptor 5 (TLR5) present in dendritic cells^41^. Chassaing et al.^43^ showed that a loss of TLR5 from DCs did not associate with inflammation or change the microbiome, but did result in a complete loss of the IL-22 immune response, which is induced by flagellin. This is in contrast to a loss of TLR5 on mouse intestinal epithelial cells, which resulted in low-grade inflammation along with metabolic syndrome and an inability to clear pathobionts^43^. Hence, They argued that this intestinal epithelial loss of TLR5 could impact the microbiome. Our results seem to align with the first scenario they describe highlighting a possible TLR5 immune response associated with DCs rather than epithelial cells, hence the observed minimal immune-response impact on the microbiome. Though one would have also expected possible competitive effect of the probiotic on the microbiome that was not observed. Further studies are required to shed more light on this result.

In support of the current literature, we found a putative effect of the probiotic, the vaccine, the prebiotic and their different permutations on the microbiome. This study is the first to describe these effects utilizing a probiotic-vectored vaccine. WT, MPER and IL1b treatments behaved similarly when comparing their beta diversity measures. These treatments resulted in a shift in the microbial community over time to a putative new equilibrium. Similarities between MPER and IL1b were also observed in the confusion matrix resulting from the random forest classification approach where the error of assignment from one of these groups to the other was comparable. While Shannon diversity highlighted a comparable reduction in diversity of these three treatments, as compared to the negative control, Chao1 highlighted a gradient decrease in richness with a clear reduction in the IL1b treatment (Figure 1). IL-1 β, part of the IL1 family, a proinflammatory cytokine with a role in modulating the adaptive immune response ^40,44^. Our results suggest that the observed richness decrease might be an outcome of the compound effect of the probiotic and the inflammatory effect of the adjuvant. An existing body of literature supports that probiotics and/or vaccination could result in modulation of the resident microbiome^45,46,47,13,18,48,49,50^. In addition, a significant decrease in richness and shift in the microbial community structure was also observed in association with the RB treatment. This treatment also stood out with zero error of assignment, utilizing the Random Forest approach, as observed for the cecal samples. These results could be attributed to the compound effect of the probiotic vaccine and the prebiotic rice bran in this case. Modulation of the microbiome utilizing rice bran^51,11^ and the effects of diet are well supported^52,53,54^. Taken together these results highlight the opportunity for synbiotics, combining probiotics and prebiotics to improve efficacy of vaccination. Careful consideration of the constitutive adjuvants and the use of diet supplementation is needed when the target is modulation of the microbiome to improve efficacy of vaccination or maintenance of the resident microbiome.

The murine microbiome was mostly dominated by taxa belonging to the Firmicutes and Bacteroidetes phyla^4^. The discriminating taxa in both fecal and cecal samples were similar, though the small cecal sample sizes blurred the impact of these taxa. In both cases, taxa were mostly dominated by Firmicutes and belonged to the Lachnospiraceae family. This family belongs to the order Clostridiales with all known taxa being strictly anaerobic and found mainly in the mammalian gut^55^. The Actinobacteria phylum was less dominant in these samples and included an OTU that belonged to the Bifidobacterium genus. This OTU had the highest impact on discriminating between the treatment levels in the fecal samples. This was not a surprise though unlike in previous research, it was not its dominance in the rice bran treatment that caused it to have a prominent effect but its presence in the negative control. Bifidobacterium is also commonly found as part of the mammalian microbiomes^56^. In mammals and humans this genus dominates in early life neonates though still comprises 25% of the microbial population within the healthy gut of adult humans^57^. No taxa of the genus Lactobacillus were observed as part of the top 30 impactful taxa in the fecal samples and only one was observed in the cecal samples, albeit all taxa had similar weight in discriminating between the treatment levels based on these samples. A more targeted approach might be required to verify the possible impact of vaccination on these specific taxa (see below).

Total IgA also showed different trends between the applied treatments. Induction of MPER-specific IgA was observed in association with the treatments that included the vaccine, which was expected. Non-parametric correlation analysis, utilizing the Spearman correlation coefficient, between the diversity measures and between the abundance of the 30 most impactful taxa and total and antigen-specific IgA did not produce highly significant associations. This indicates that either other immunological factors, in addition to the direct impact of the vaccine vector, were at play or that a more targeted approach is required to better assess these associations. A possible more direct approach to assess the correlation between the change in IgA and its possible microbiome targets in the gut is to focus on the study of the IgA-coated microbiota. These taxa represent the putative resident microbiome most affected by changes in the immune response. Changes in abundance and presence/absence of components of this part of the microbiome could highlight possible off-target effects of the vaccine whereby resident taxa that are evolutionarily or functionally similar to the vaccine vector might be targeted by the immune response. We are currently assessing this hypothesis in a proof of principle follow-up study.

## Materials and Methods

### Study Design and Vaccine Strains

6-8 week old BALB/c mice (Jackson Laboratory) were housed together in isolated cages based on vaccine strain to be received. Animal gut microbiomes were normalized using multiple procedures including standardized diet, identical cage locations, cage swapping, and oral gavage of cecum content one week prior to first vaccination^58^. Six animals per group received either *Lactobacillus acidophilus* wild-type strain (WT), WT expressing the membrane proximal external region from HIV-1 within the context of the major S-layer protein A (MPER), MPER secreting interleukin-1β (IL1b) or MPER expressing surface flagellin subunit C (FliC). Five mice received MPER and a custom 10% rice bran diet (RB) from Envigo, prepared as previously described^59^. The six mice in the negative control group (NG) received only the dosing buffer composed of 8.4 mg/mL NaHCO_3_ and 20 mg/mL soybean trypsin inhibitor (SIGMA T9128) in ultrapure water^29,60^. Bacterial cells were prepared from overnight culture and suspended in dosing buffer. Each animal received 5 x10^9^ CFU/day in 200 µL of dosing buffer by oral gavage for three consecutive days at weeks 0, 2, 4, 6, 8, and 10 (six 3-day doses) as previously described^39,61,62^. Samples were collected on the first day of each dose, prior to dose administration. Rice bran diet group samples were also collected one week prior to starting the RB diet (week −1) and at week 12.

The care and use of experimental animals complied with the guidelines of Colorado State University (IACUC 14-5332A) and all experiments were performed in accordance with these guidelines and regulations. All animals were age matched, maintained in specific pathogen free conditions, individually tracked and monitored daily for clinical signs of stress or illness, including but not limited to changes in skin and hair, eyes and mucous membranes, respiratory system, circulatory system, central nervous system, salivation, diarrhea, or lethargy. Upon arrival, animals were housed socially (*n* = 2–5) in commercially available, individually ventilated caging systems with a 12 h light/12 h dark cycle. Animals were provided *ad libitum* commercial irradiated rodent chow (Teklad Global) or the rice bran diet and tap water filtered via reverse osmosis in autoclaved water bottles; all bedding and enrichment materials were autoclaved prior to use and changed regularly.

### Sample Collection

Fecal pellets were collected from mice in cleaned collection cups and transferred to pre-weighed collection tubes. Pellets for supernatant extraction were weighed and 10 µL of 2x ProteaseArrest (G-Biosciences) was freshly diluted in cold PBS (from 50x stock) per mg of fecal pellet weight within 5 minutes of excretion. Samples were then homogenized via three 20-second cycles (4 m/s) using a FastPrep-24 Homogenizer (MP Biomedicals) and centrifuged at 10,000 RCF, 4°C for 10 minutes. The supernatant was removed and stored at - 80°C for later analysis. Fecal pellets for microbiome analysis were collected directly from the anus into collection tubes, immediately placed on ice, and stored at −80°C until extraction. Cecal contents for microbiome analysis were obtained by necropsy at week 10 for treatments NG, WT, MPER, IL1b, and FliC, and at week 12 for RB, squeezed into collection tubes, immediately placed on ice, and stored at −80°C until extraction. Animals were euthanized by CO_2_ asphyxiation and subsequent cervical dislocation, according to IACUC protocol.

### ELISA

MaxiSorp high-binding 96-well plates (Nunc) were coated overnight at 4°C with 100 µL/well of 10 µg/mL goat anti-mouse IgA antibody (Bethyl Laboratories, Inc.) for total IgA ELISA, or with 1 µg/ml of synthetic 17-mer HIV-MPER peptide (GNEQELLELDKWASLWN, Bio-Synthesis Inc.) for antigen-specific ELISA, both suspended in carbonate coating buffer (15 mM Na_2_CO_3_, 35 mM NaHCO_3_ in ultrapure water). Wells were blocked with 1% BSA in PBS for 1 hour at room temperature. After washing (0.05% Tween 20 in PBS), fecal supernatant was serially diluted 1:2 (from 1:10 to 1:640) in PBS with 1% BSA and 0.05% Tween 20, added to wells, and incubated for 2 hours at room temperature. Following washing, HRP-conjugated goat anti-mouse IgA antibody (50 ng/mL, Bethyl) was added and incubated for 1 hour at room temperature. Color development with 3,3’,5,5’-tetramethylbenzidine (TMB) was terminated after 15 minutes with sulfuric acid and absorbance (570-450 nm) was measured.

To quantify total IgA, 1:10 dilutions of Mouse Reference Serum (Bethyl Laboratories) starting at 1000 ng/mL were run on each plate in duplicate to generate a standard curve. Fecal supernatants from each animal’s week 0 time point were included in the assay to determine endpoint titer for antigen-specific IgA. The optical density cutoff was calculated as the mean value of all negative controls for each vaccination group + 3.365 standard deviations, based on the 99% confidence interval standard deviation cutoff multiplier for n=6^63^.

### DNA Extraction and Sequencing

Microbial genomic DNA was extracted from whole fecal pellets using PowerFecal DNA Isolation Kit (QIAGEN); buffer-only controls were included in each extraction batch. The hyper variable region 4 (V4) of the 16S rDNA gene was amplified by PCR according to the Human Microbiome Project protocols^64,65^. Primers used for multiplexed 16S library generation included an Illumina adapter, an 8-nt index sequence for barcoding, a pad and linker, and the 16S-specific sequence, as described previously^65^. ZymoBIOMICS™ D6305 Microbial Community Standard (mock community) and no-template controls were included in each PCR plate. Dual-indexed library molecules were then purified using Sera-Mag beads (GE Life Sciences) to select for DNA fragments larger than 150bp. Purified library molecules were quantified using Quant-iT Broad-Range dsDNA kit (Invitrogen) according to manufacturer’s protocols. Libraries were pooled at equimolar concentrations and quality controlled prior to sequencing on an Illumina MiSeq at the Colorado State University’s Next Generation Sequencing Core Facility. Paired-end 2×250 reads and index reads for multiplexing were generated as previously described^65^. Samples were sequenced targeting an average read-depth of 4×10^4^ reads per sample. All raw sequence data were deposited and are available on the National Center for Biotechnology Information’s (NCBI) Sequence Read Archive (SRA) repository under accession number PRJNA542488.

### Data Processing and Bioinformatics

Sequence data were processed using mothur^66^ version 1.39.5 and utilizing the developers’ standard operating procedure (SOP) for OTU calling and taxonomic classification of MiSeq data first presented in Kozich, et al., 2013^65^. For alignment and classification within this SOP we used the SILVA database^67^ version 128. Sequencing error rate was assessed using the ZymoBIOMICS™ microbial community standard (mock community) described above, which includes 8 bacterial and 2 fungal species. Our sequencing included five mock community samples, nine negative controls with no sample, and four negative controls with no template. Clustering, for OTU identification, was performed using OptiClust^68^ utilizing 0.97 sequence similarity. We used maximum counts per OTU of all mock samples to identify a cutoff number of reads per OTU to guard against overestimation of sample diversity. Rarefaction curves were generated using the package vegan^69^ as implemented in R version 3.4.4^70^ to assess diversity and suitability of depth of coverage per sample. The resulting OTU table was utilized in further data analyses as follows. Results of data processing and the associated bioinformatics can be found in the Supplementary File 1.

### Diversity and IgA analysis

#### Alpha Diversity

Chao1 and observed richness along with Shannon and inverse Simpson diversity indices were computed utilizing the R package phyloseq^71^. We used a Bayesian framework to fit and compare 30 putative models that evaluate linear and nonlinear associations between time and these measures of diversity. Model fitting was done using the R-package rjags^72^ an interface to the Just Another Gibbs Sampler (JAGS) software^73^ and model selection was done using the Deviance Information Criterion (DIC)^74^ also implemented in rjags. Supplementary File 2 provides detailed description of the general characteristics of the different models used. The model with minimum DIC was selected to be the best-fit model. Convergence of the Markov Chain Monte Carlo (MCMC) sampling was assessed using the trace diagrams associated with the chosen models and Gelman and Rubin’s convergence diagnostic^75^ based on three chains that were run for 500,000 iterations using the first 200,000 as burn-in and sampling every 500 steps to assure sample independence. This resulted in 600 samples per chain (a total of 1,800 samples) that we used to construct the posterior distribution of the parameters. Times −1 and 12 associated with the RB treatment were dropped from this analysis to put all treatments on the same footing.

#### Beta Diversity

Nonmetric Multidimensional Scaling (NMDS)^76^ was used on the OTU level to assess possible trends and clustering in the microbial community structure per treatment condition and per time point. NMDS was performed separately per treatment level using the vegan package and utilizing Bray-Curtis dissimilarity^76^ and was based on data transformed using Cumulative Sum Scaling^77^.

#### IgA

Total IgA was compared between treatment levels, other than RB, using the same Bayesian framework described above for alpha diversity. Total IgA data were not collected for the RB treatment. MPER-specific IgA data were transformed by first replacing zero measurements with 1 and then computing the log. This was done to overcome the high variation observed in the measurements.

### Trends and Association on the Taxonomic Level

#### Bar graphs

Utilizing relative abundance data based on the resulting OTU table, bar graphs were generated using the ggplot2^78^ package in R. These plots were generated for the relative abundance data at the phylum level and meant to describe the microbial community structure per time point under each of the treatments.

#### Random Forest Classification

We used Random Forests (RF)^79^ to identify the major microbial taxa that influenced the accurate prediction of the state of vaccination (treatments) over the period of the experiment. We used an iterative approach to identify the optimal number of features (variables) to incorporate in constructing the regression trees in the RF. In this approach we iteratively fit a random forest including one all through 100 features using the mtryStart option in the function tuneRF in the randomForest^80^ package. This was done 100 times and the out-of-bag (OOB) error was averaged over all runs. The number of features to include in sampling to construct the regression trees in the RF was chosen to be that with the minimum median OOB. We used the parameter ntree = 2000 for the total number of trees to grow in the forest and set the parameter importance to TRUE to assess importance of the different features in prediction. The predictive features included all OTUs in addition to time of vaccine administration per treatment level to account for the impact of possible change over time.

#### Association with Immune Response

We used Spearman’s correlation coefficient^81^ to assess the association between time, total IgA, antigen-specific IgA and the alpha diversity measures. This was also done to study the associations between taxa identified as most predictive using RF and the immune response.

## Author contributions

ZA, JL and GAD conceived the ideas and developed the experimental design of this project, ZA also performed the analyses and wrote the manuscript, JL performed the experiments associated with the immunology and participated in the writing of the manuscript, AL also contributed to the immunological experiments and lead the microbiome sequencing and contributed to the writing of the manuscript, BE also contributed to the sequencing portion and the to the writing of this manuscript, finally, ER also contributed to the design of the experiment and to the writing of this manuscript.

## Data and Materials availability

Raw sequence data are available from the National Center for Biotechnology Information’s (NCBI) Sequence Read Archive (SRA) under accession number PRJNA542488.

## Supplementary File 1: Results of data processing and bioinformatics

### Data Processing and Bioinformatics

Using mothur-MiSeq data processing SOP (described in *Materials and Methods*) we identified 1,405 putative OTUs (range 100-249 OTUs per sample) prior to data filtering. Sequencing depth per sample ranged between 8,592 and 115,713 reads. The sequencing error rate was calculated to be 2.414×10^−7^ with 89 identified putative OTUs within the mock community samples. Given that the average number of reads associated with the mock community samples was 33,505 and allowing for an error of read-OTU misidentification of 1 in 1000 reads we set a cutoff of 4 (rounding up from 3.3) as the minimum number of reads per OTU required to consider that OTU to be correctly identified. This cutoff resulted in a reduction in the number of OTUs identified in association with the mock community to 15 from 89 (true total is eight). Six of the erroneously identified OTUs belonged in low abundance to only one of the five mock community samples. This also restricted the number of taxa identified as present in the negative control with no sample to four with number of reads less than or equal to 10 in five of these samples and in the negative control with no template to two OTUs with number of reads less than or equal to 4 in three of the four samples. This cutoff was subtracted from all OTU counts per sample resulting in a conservative, total number of putative OTUs of 271 (range 53-165 OTUs per sample) in 245 samples associated with the above described experimental design (including both fecal and cecal samples). Figure S1 shows the resulting rarefaction curves for the fecal and cecal samples, separately, and indicates adequate depth of sequencing to capture sample diversity.

In further assessing the quality of the outcome data we generated NMDS plots separately per treatment level to evaluate time trends and possible outliers. This preliminary data exploration resulted in identification of all samples associated with the positive control (WT) at the second time point as being outliers. These samples grouped together but were quite distant from all other samples belonging to all other time points within that treatment. In further investigating these samples we concluded that they were sampled directly after introduction of the probiotic at that time point resulting in over-dominance of the lactobacillus genera (data not shown). This deviates from our sampling procedure used through the experiment (described above); hence, these samples were dropped from further analyses.

## Supplementary File 2: Univeriate Models for Analysis of Diversity and Total-IgA Data

### Rationale

The experimental design involved in this project includes six treatment levels (WT, MPER, IL1b, FliC, RB and NG) as described in the methods. We consider these treatment levels to be fixed effects that we intend to compare. Nested within these treatment levels we have 6 mice (5 for RB) sampled over 10 time points. Accordingly, we have a repeated measurement/time series design with mice taken as random effects. We evaluate a linear, quadratic and cubic time trend. In what follows we focus on Alpha diversity measures specifically Chao1 richness and Shannon diversity; we use similar models for both and for Total-IgA. For any model that accounts for trend associated with time we include all lower terms in that model when a higher term is included. For example, models that include a cubic term will also include both the quadratic and linear terms. Those models that include a quadratic term will also include the linear one. We use the Deviance Information Criaterlion (DIC) to assess and identify the best models that fit the data as described in the *Methods*. In the case of the cecal samples, where there is no trend over time, we only compare the treatment effects.

### Model for trends in alpha diversity

Let *y_itj_* be the observed diversity measure for mouse *j* at time *t* nested within treatment *i*. Following is the full model that includes all time trends assuming that each mouse *j* nested within each treatment *i* (shown as *i*[*j*]) follows its own separate cubic time trend:

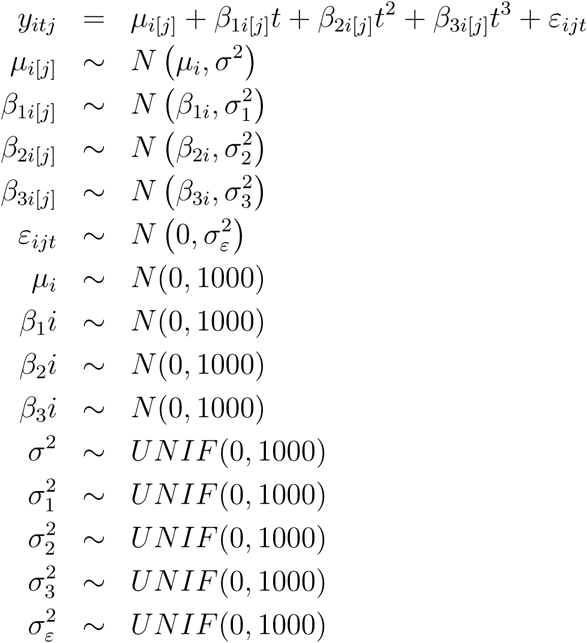

Where *N* (*m, n*) represents a normal distribution with mean *m* and variance *n*, *UNIF* (*a, b*) represents a uniform distribution with boundaries *a* and *b*, *µ_i_*_[*j*]_ is the intercept associated with each mouse *j* nested within treatment *i*, *β*_1*i*[*j*]_, *β*_2*i*[*j*]_ and *β*_3*i*[*j*]_ are the coefficients associated with the linear, quadratic and cubic trends per mouse *j* nested within treatment *i*. *µ_i_*, *β*_1*i*_, *β*_2*i*_ and *β*_3*i*_ are the intrecept, linear coefficient, quadrtatic coefficient and cubic coefficient, repsectively, associated with treatment *i* and these are the parameters of intrest and have been used to plot the trends in the figures. the *σ*’s repesent the variances.

The model above could be resuced in 29 different ways (resulting in 30 models to compare) that restrict the trend to quadratic or linear and that restrict the model to represent the treatments rather than accounting for the individual random effects of each mouse. For eaxample, a model that accounts for quadratic trend per treatment level is as follows,

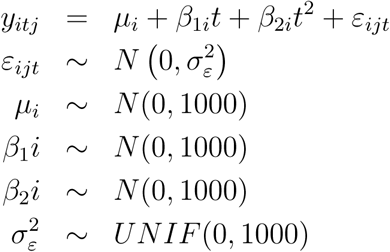

A model that assumes a linear trend shared for all treatments with different intrecept per treatment is as follows,

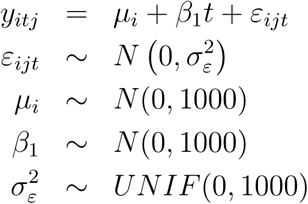

A model assuming no trend associated with time but with mouse specific intercepts is as follows,

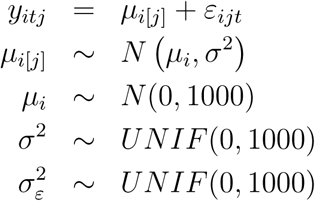

A model that assumes no trend but with intercepts associated with the treatment only (no effect of the mouse) is as follows,

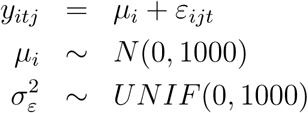

The last two models along with the following simplest null model were used in association with the cecal samples as well as the fecal ones. The minimal null model is as follows,

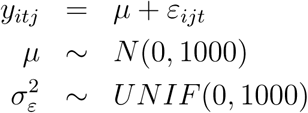

With generalized mean and variance representing no differences between the mice and treatments and no observed time-trend.

## Supplementary Figures

**Figure S1:**
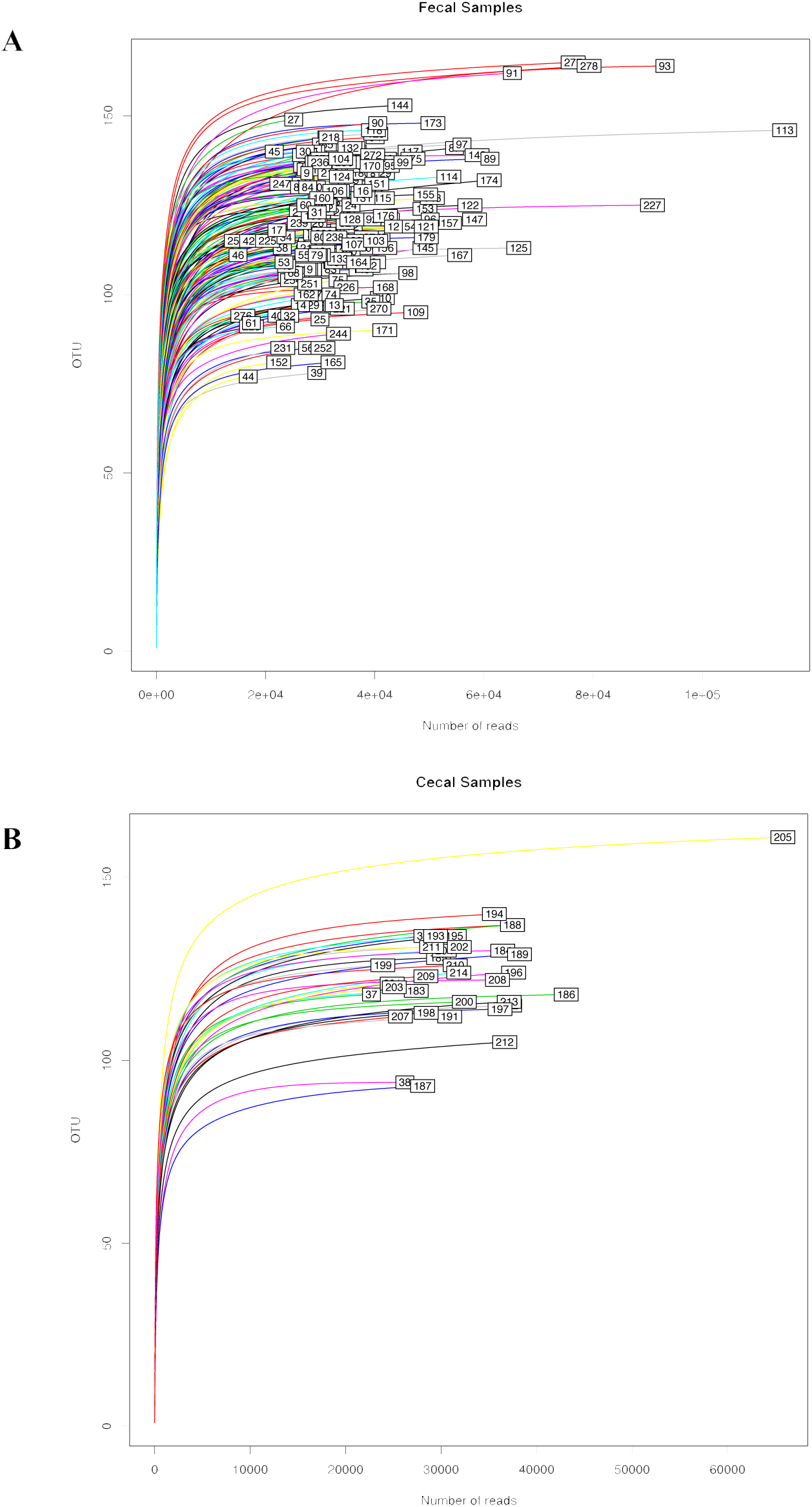
Rarefaction curves describing depth of sequence coverage for both (A) fecal and (B) cecal samples.

**Figure S2:**
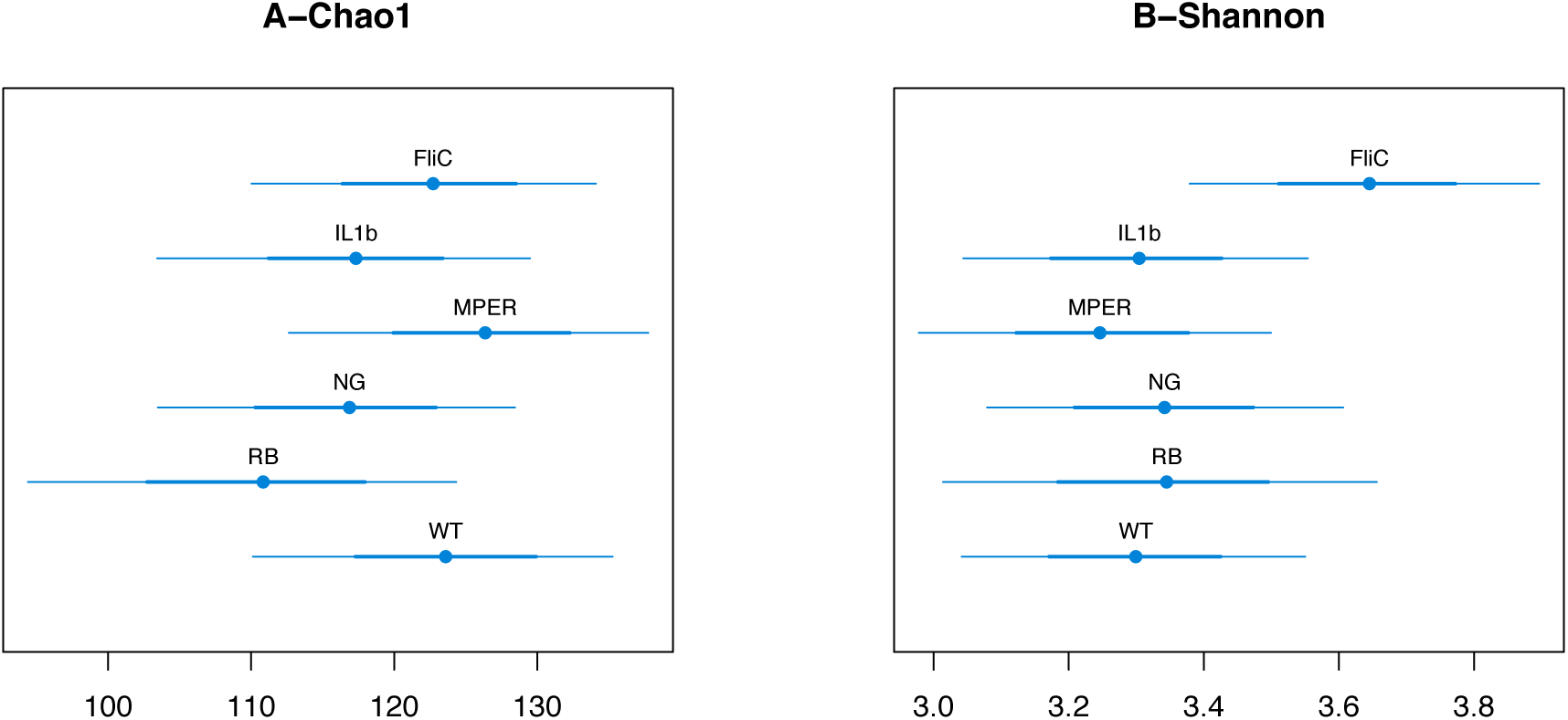
(A) 95% credibility intervals of the expected Chao1 richness treatment levels for cecal samples. (B) 95% credibility intervals of the expected Shannon diversity index per treatment levels for cecal samples.

**Figure S3:**
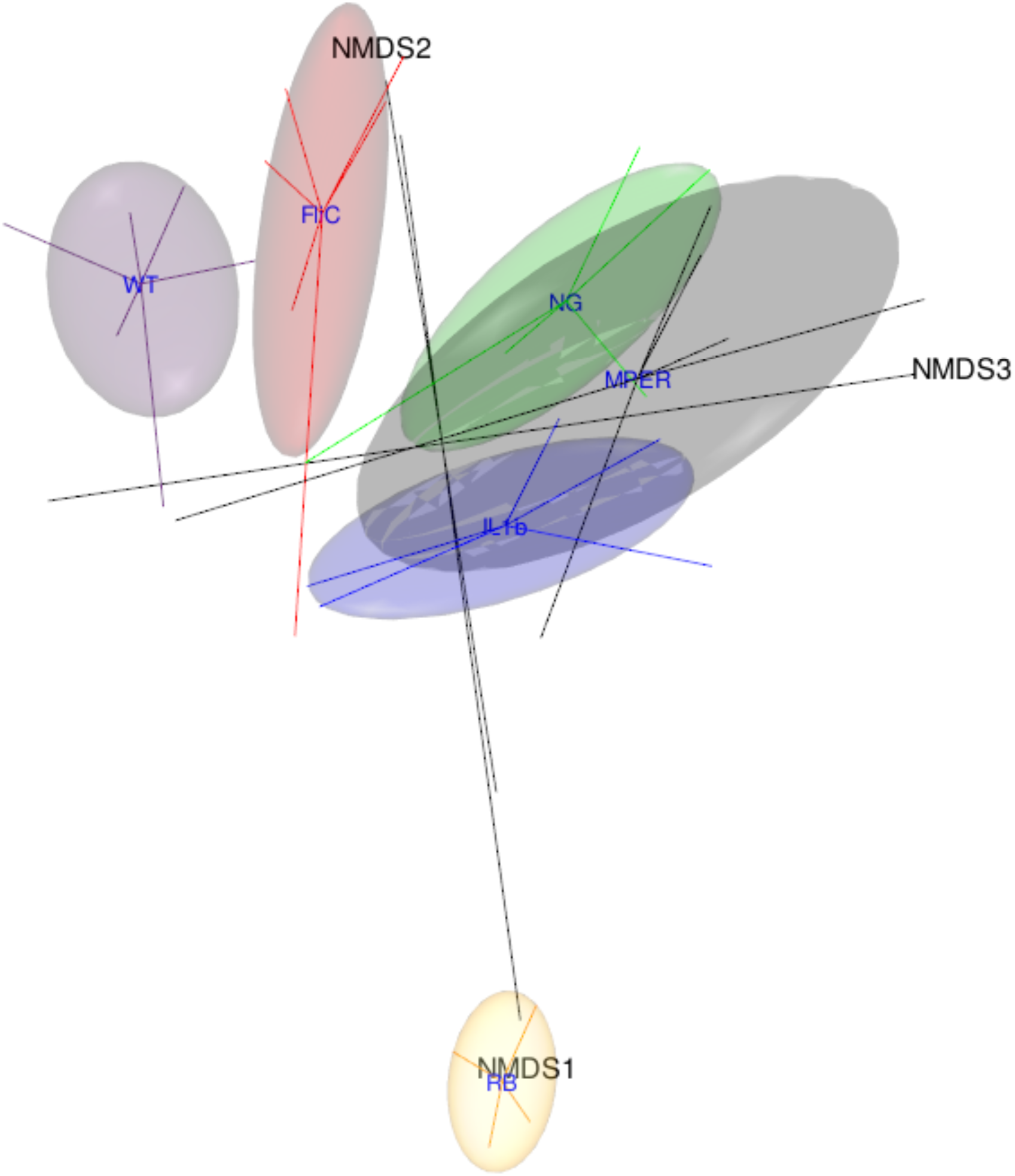
Three-dimensional (3D) nonmetric multidimensional scaling (NMDS) ordination plot of the beta diversity of cecal samples measured using the Bray-Curtis distance and aggregated by treatment level. The figure highlights the different data points as tips of the star shapes emitted from the centroids representing the treatment levels and the associated 95% confidence ellipsoids.

**Figure S4:**
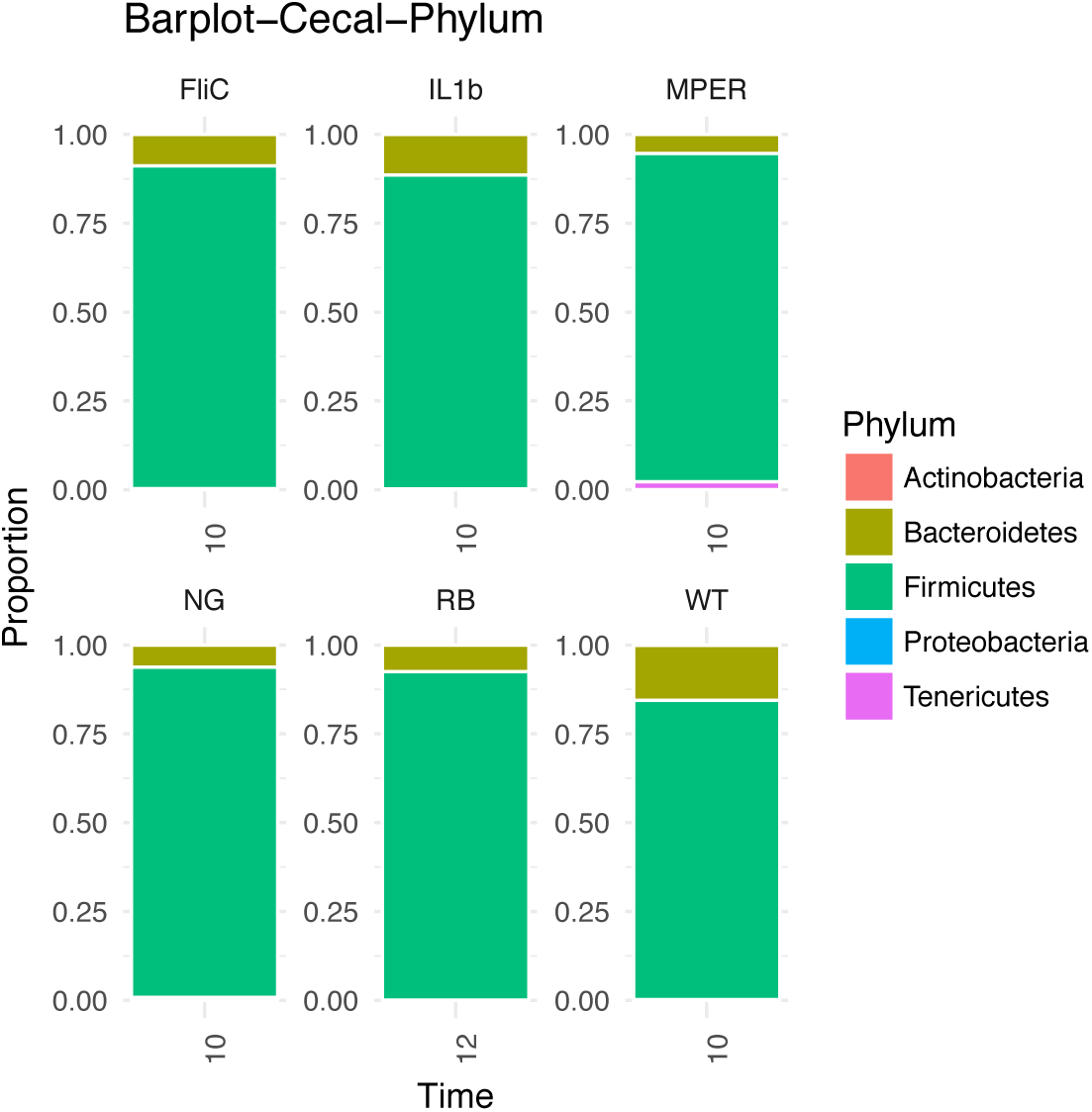
Bar plots representing the phylum level taxonomic distribution per treatment level per time point of the cecal samples.

**Figure S5:**
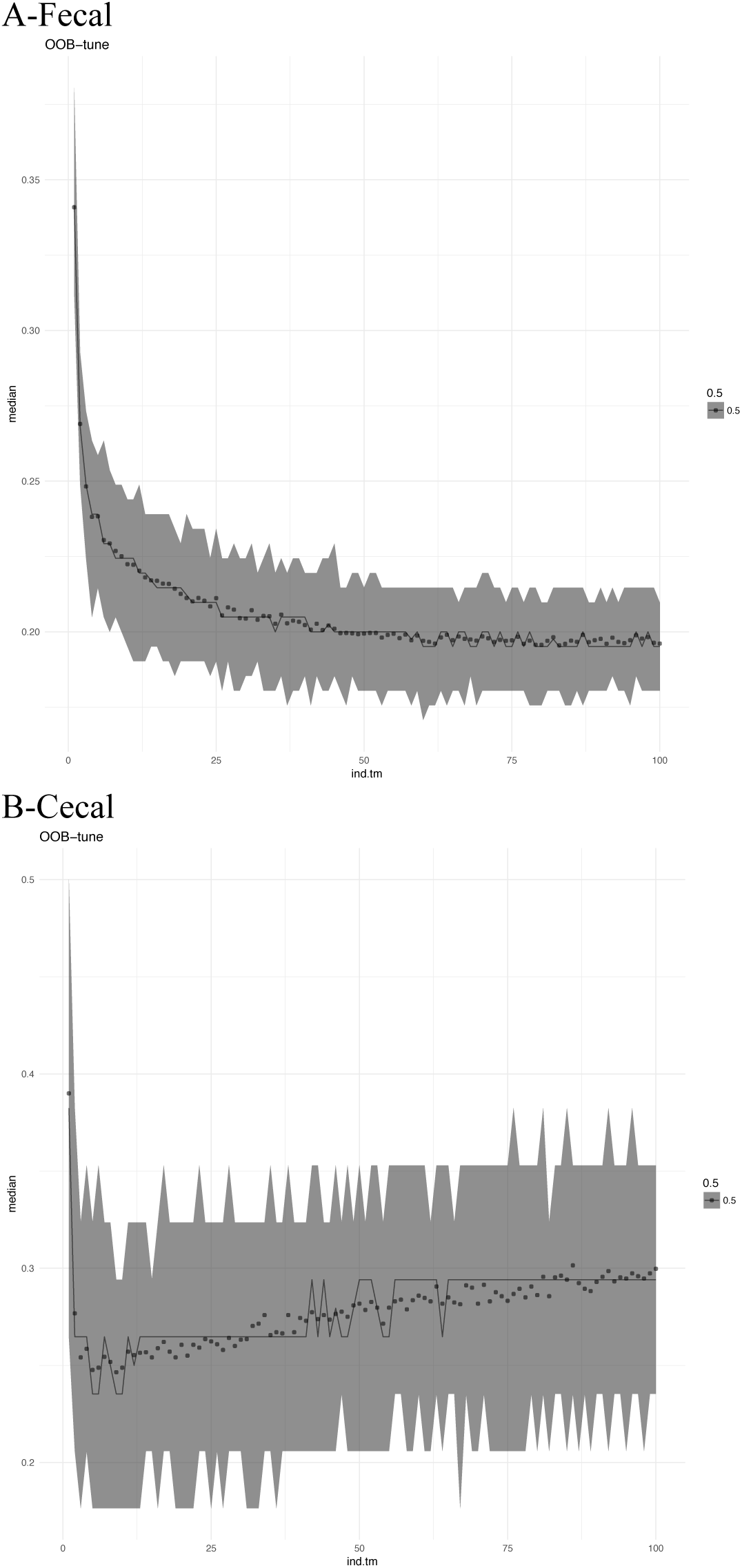
The out of bag (OOB) tuning plots used to select the optimal number of variables to use in the random forest algorithm. First minimum OOB for fecal samples was observed when using 53 variables and 9 for cecal samples.

**Figure S6:**
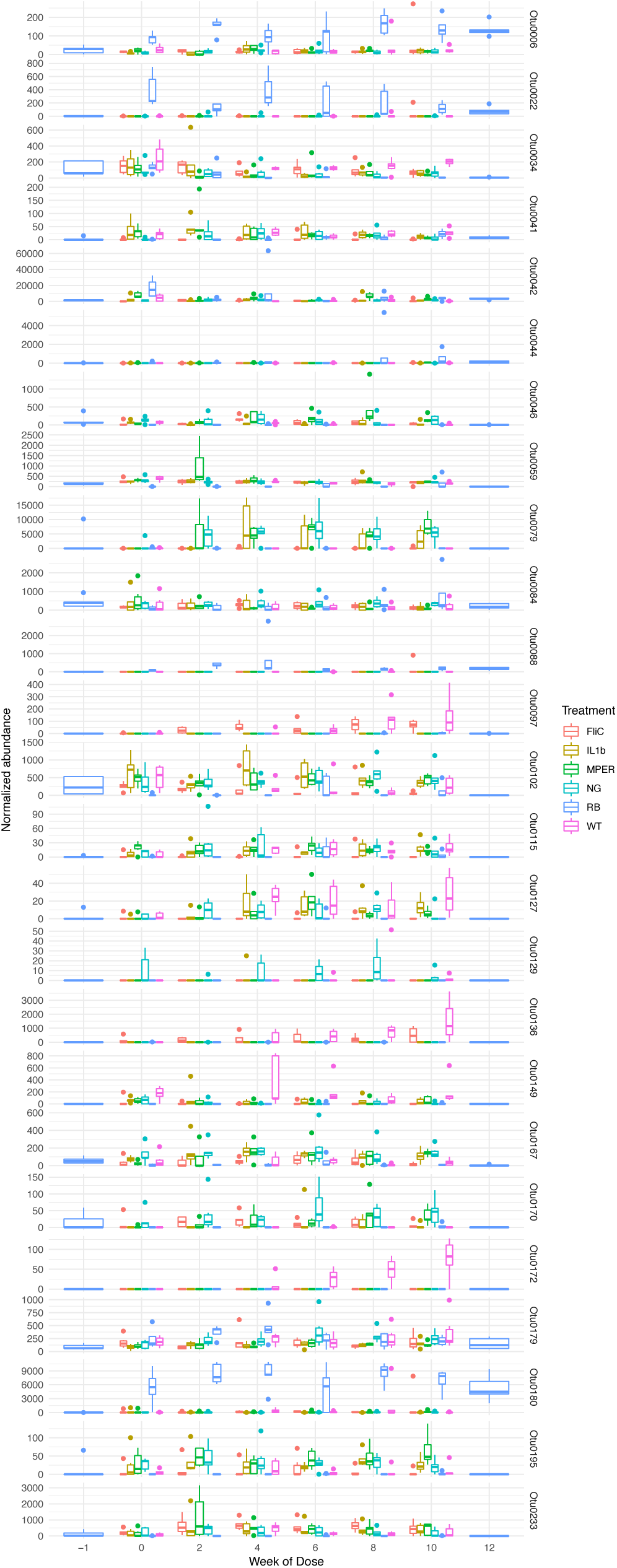
The normalized abundance of the 25 most impactful OTUs, after the five introduce in Figure 6, associated with the fecal samples.

**Figure S7:**
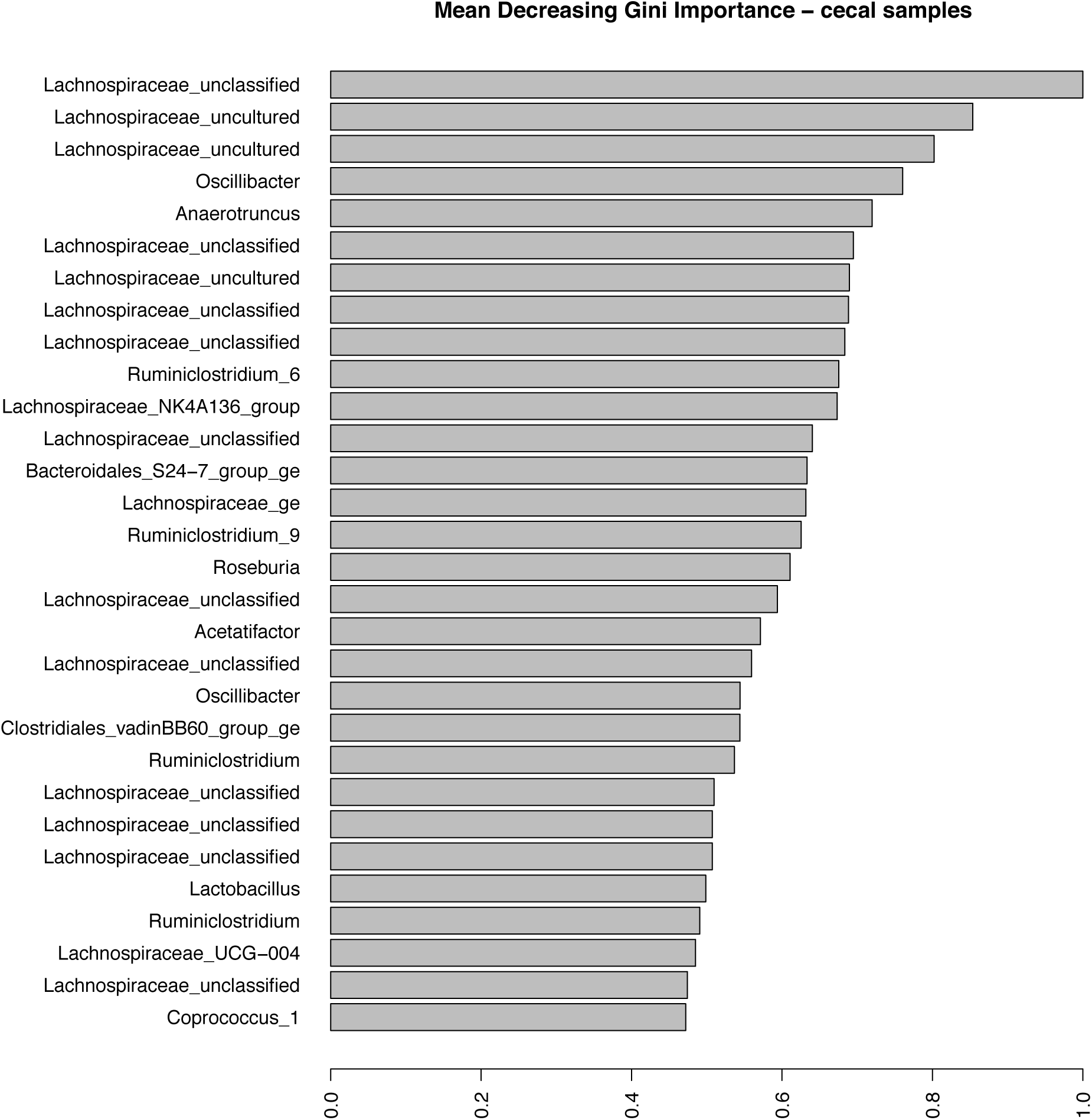
The Mean Decreasing Gini OTU importance plot for the cecal samples. X-axis represents the Gini importance measure where high values represent high impact of the OTUs presented on the y-axis.

**Figure S8:**
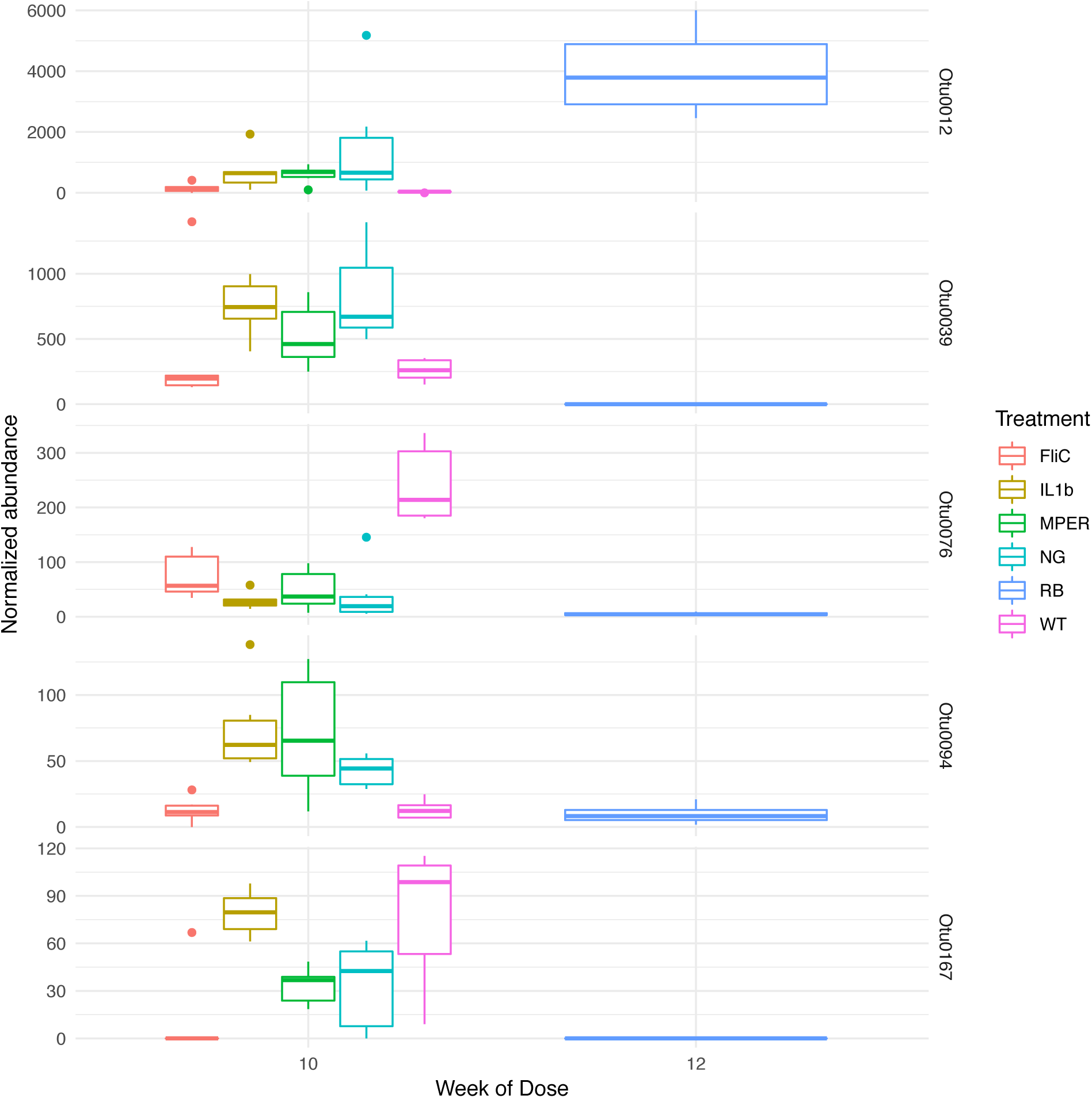
The normalized abundance of the five most impactful OTUs associated with the cecal samples as observed over time: OTU0076, OTU0094 and OTU0039 were unclassified or uncultured Lachnospiraceae; OTU0012 belonged to the Lachnospiraceae_NK4A136_group genus; and OTU0167 belonged to the Oscillibacter genus.

**Figure S9:**
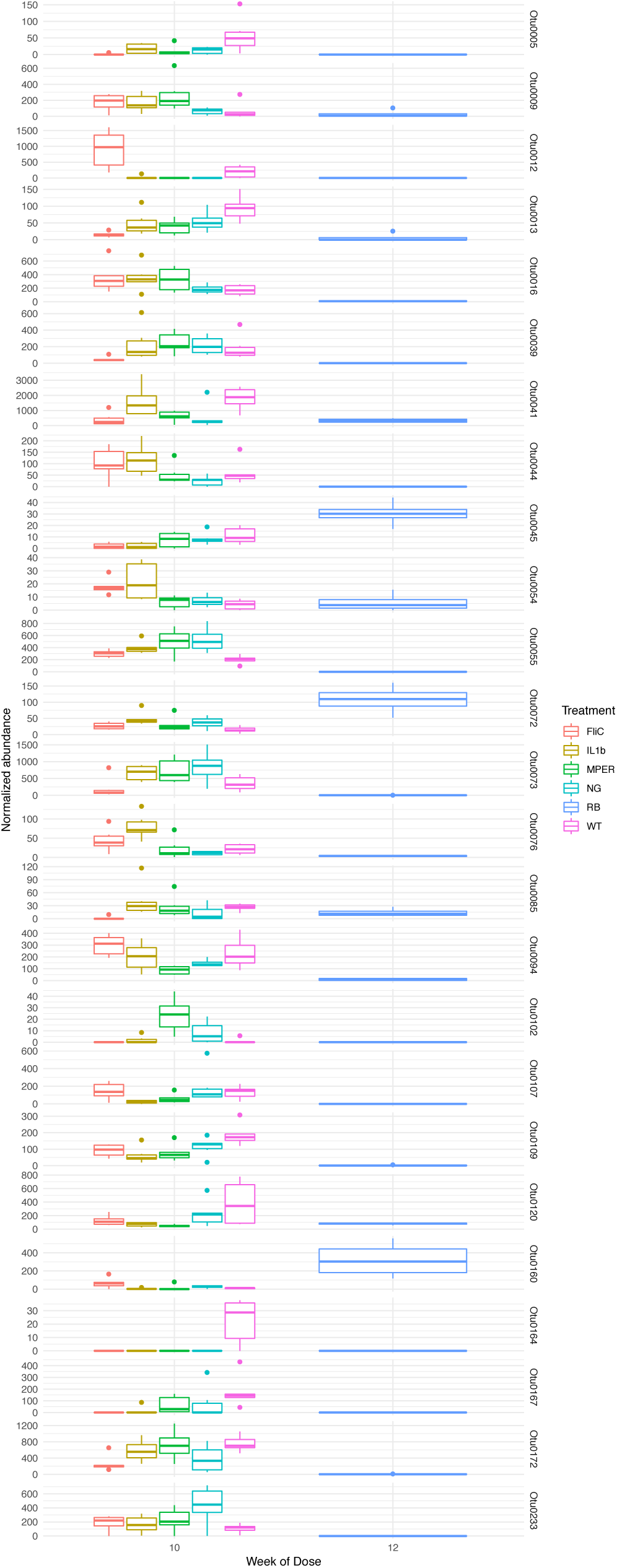
The normalized abundance of the 25 most impactful OTUs, after the five introduce in Figure S8, associated with the cecal samples.

**Figure S10:**
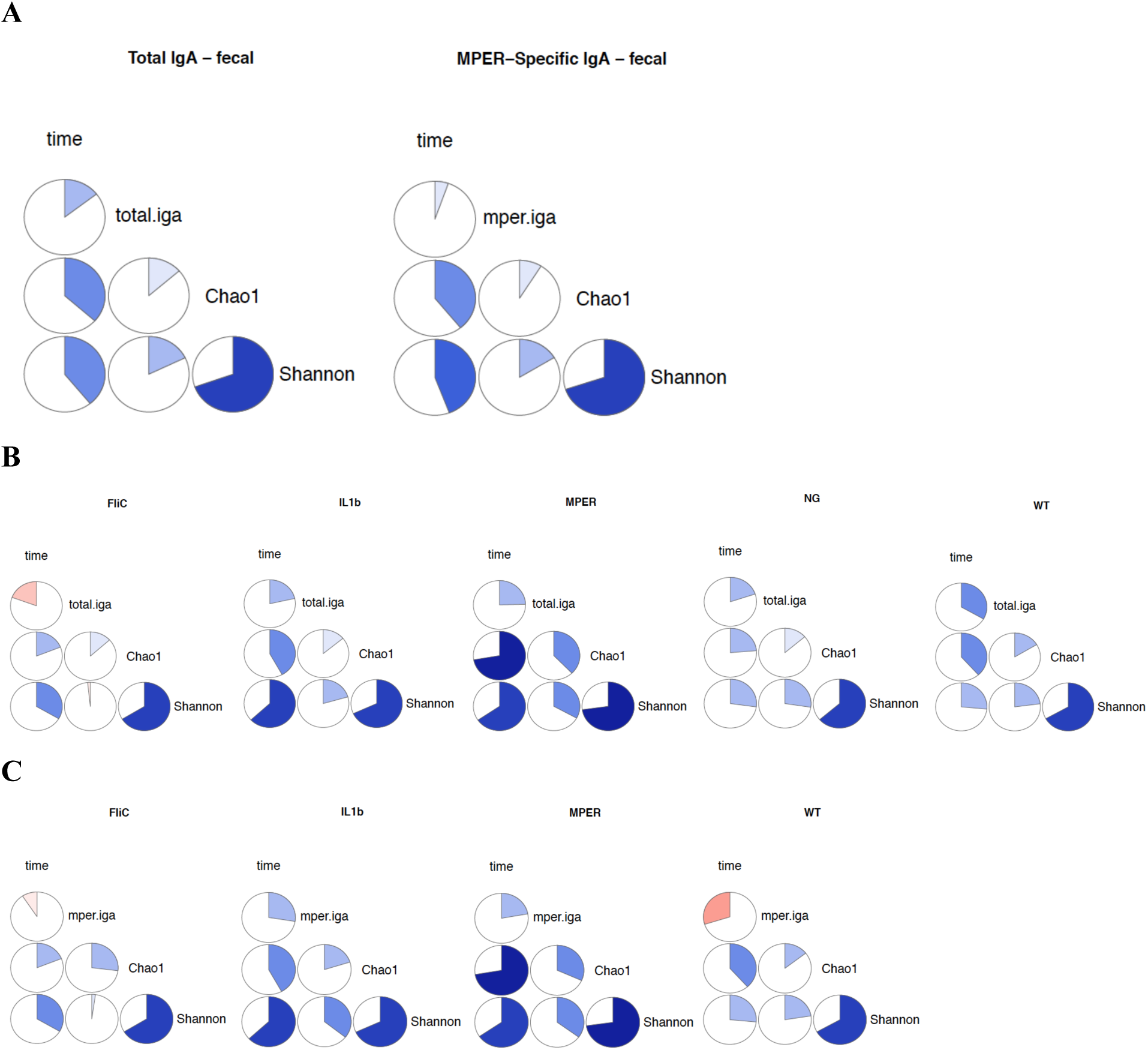
Pie charts representing Spearman correlation between the Chao1 richness, Shannon diversity, time and total IgA and MPER-specific IgA, separately, for both the (A) combined data and per treatment for (B) total-IgA and (C) MPER-specific samples.

**Figure S11:**
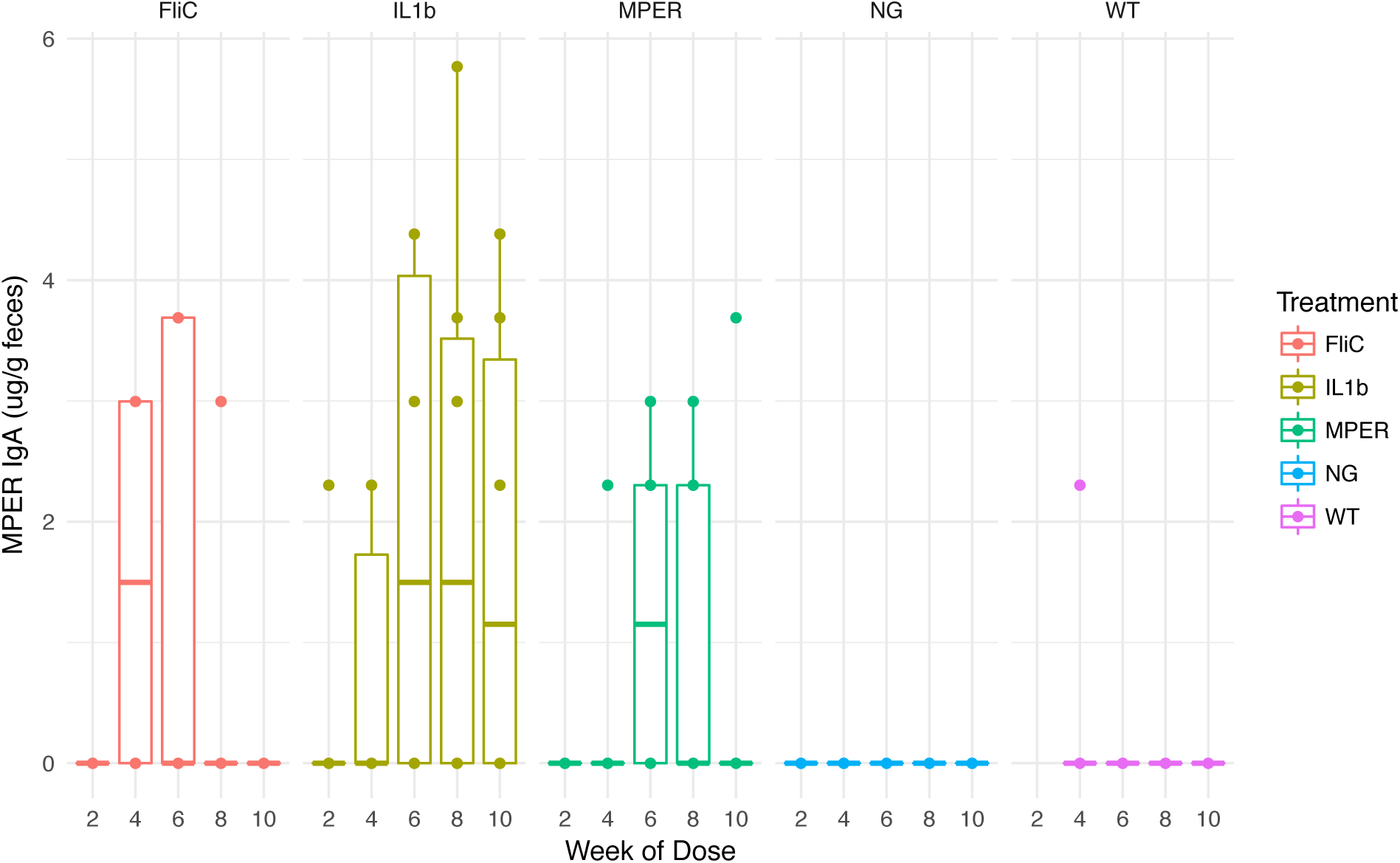
Box and whisker plots the MPER-specific IgA per time point (0, 2, 4, 6, 8 and 10) under each treatment level. Centerlines represent the median MPER-specific IgA per time point.

**Figure S12:**
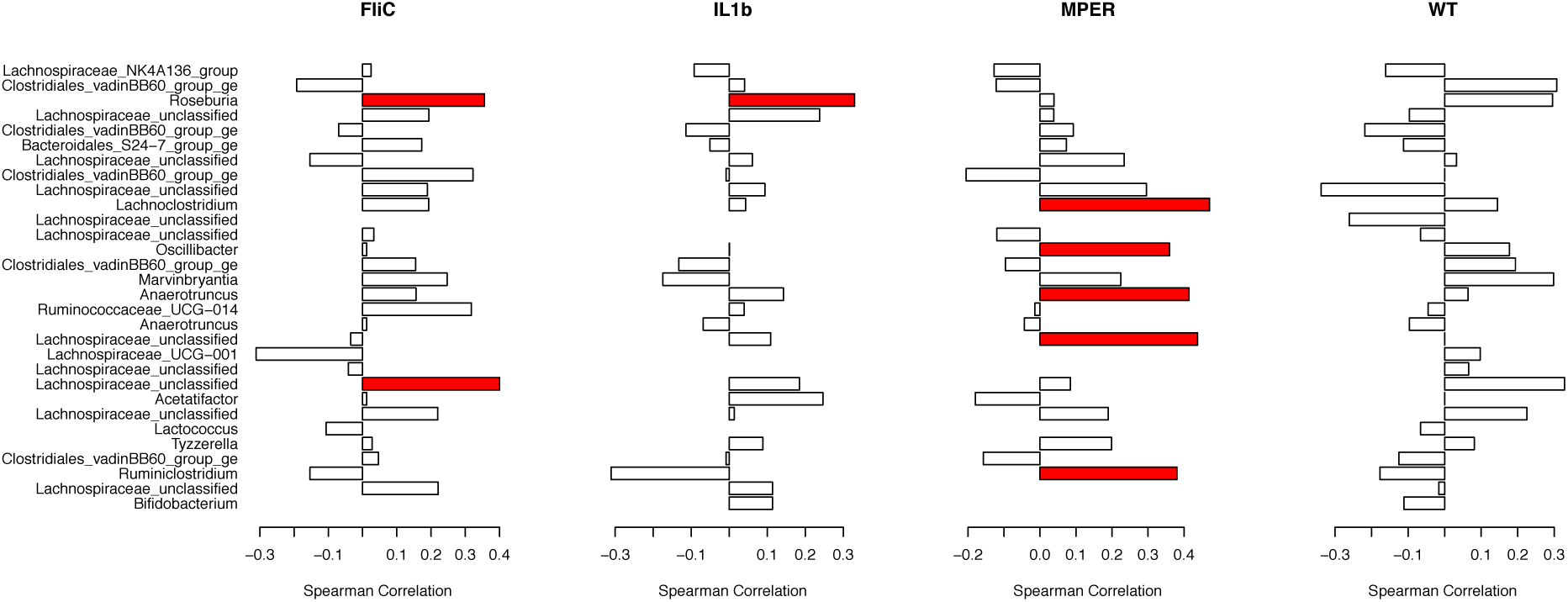
Spearman correlation plots linking the 30 most impactful taxa observed with the fecal samples and MPER-specific IgA per treatment. Red and blue represent significant positive and negative correlations, respectively, at the 0.1 level of significance with no correction for multiple testing.

## Supplementary Tables

**Table S1:** DIC-optimal models for Chao1 richness and Shannon diversity. *Y_k(i)t_* is the observed diversity for mouse *k* nested within treatment *i* at time *t*. *µ* represents the intercept, *β_1_* is the slope for the linear time component *t* and *β_2_* is the slope for the second order time component *t^2^*. *ε* is the error term. Further model details are in the Supplementary File 2.

|  |  |  |
| --- | --- | --- |
| Alpha Diversity | DIC Optimal Model |  |
| Chao1 | $Y_{k(i)t} = \mu_{k(i)} + \beta_{1k(i)}t + \beta_{2k(i)}t^2 + \varepsilon_{k(i)t}$ | |
| Shannon | $Y_{k(i)t} = \mu_{k(i)} + \beta_1t + \varepsilon_{i(k)t}$ | |

**Table S2:**
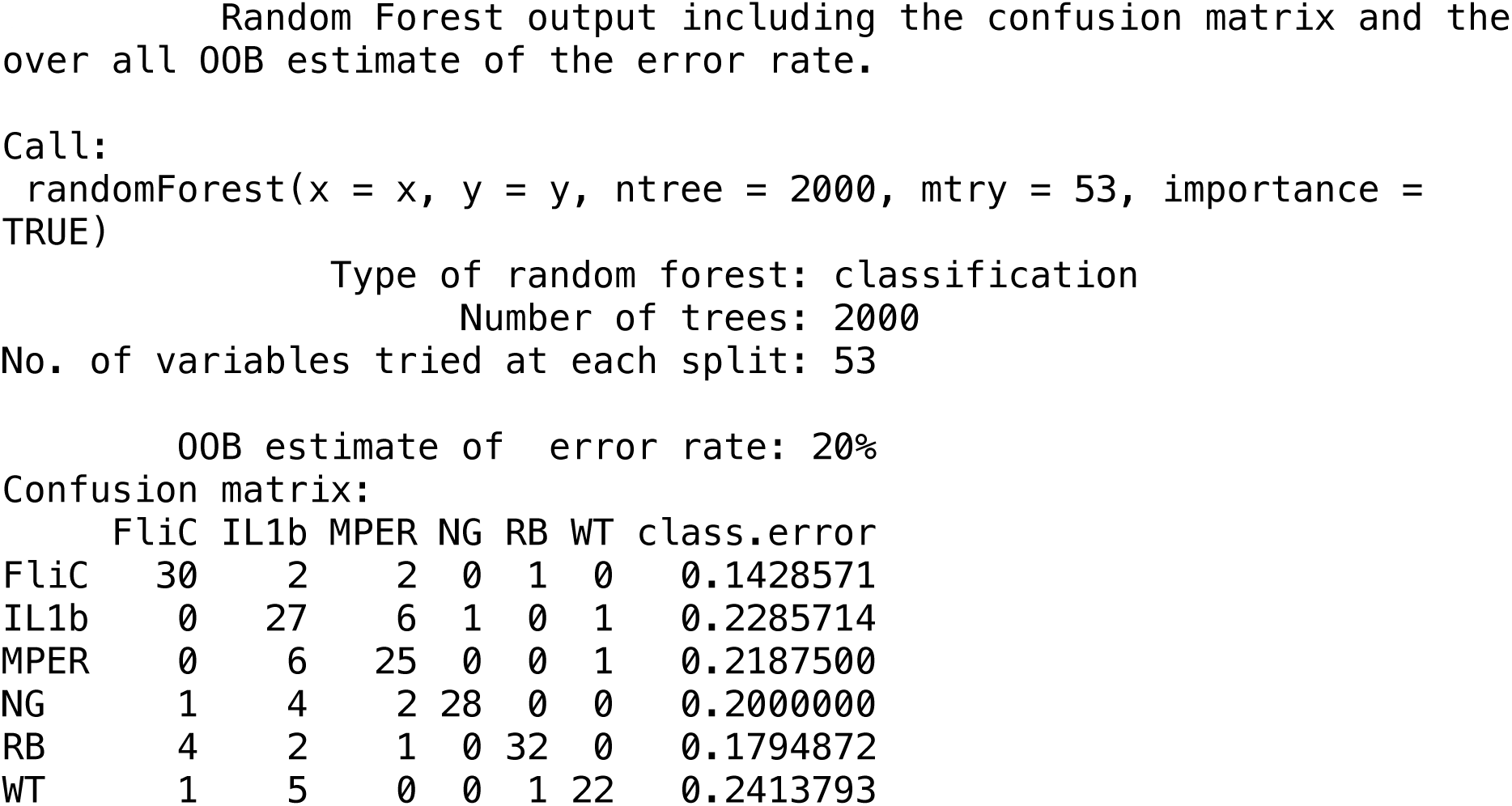
Confusion Matrix and out of bag (OOB) error rate of Random Forest classification of the fecal samples. Numbers on the diagonal of the table correspond to correctly classified samples while the off diagonal numbers represent misclassified samples. Rate of misclassification per class (treatment) is captured in the “class.error” column.

**Table S3:**
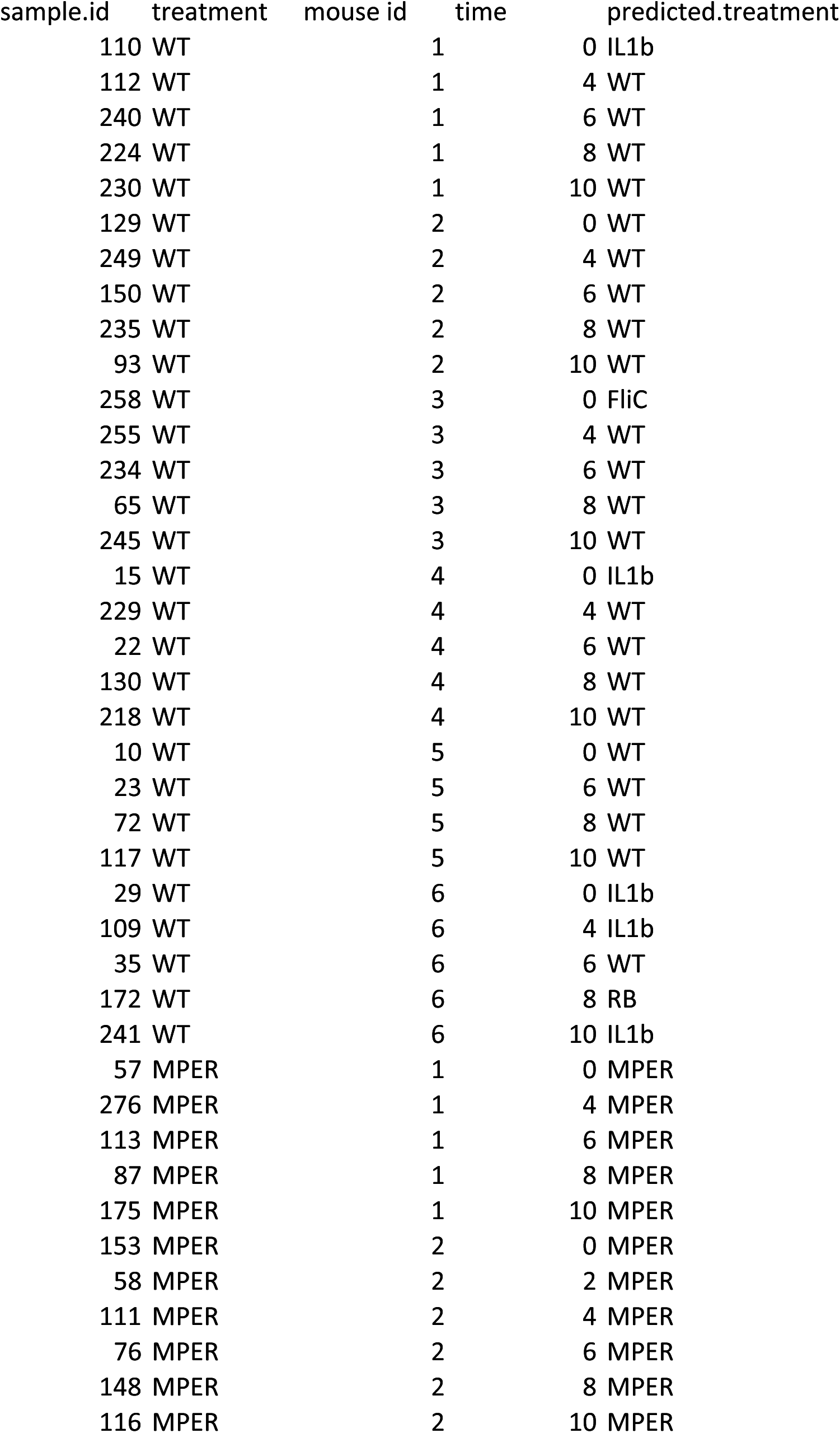
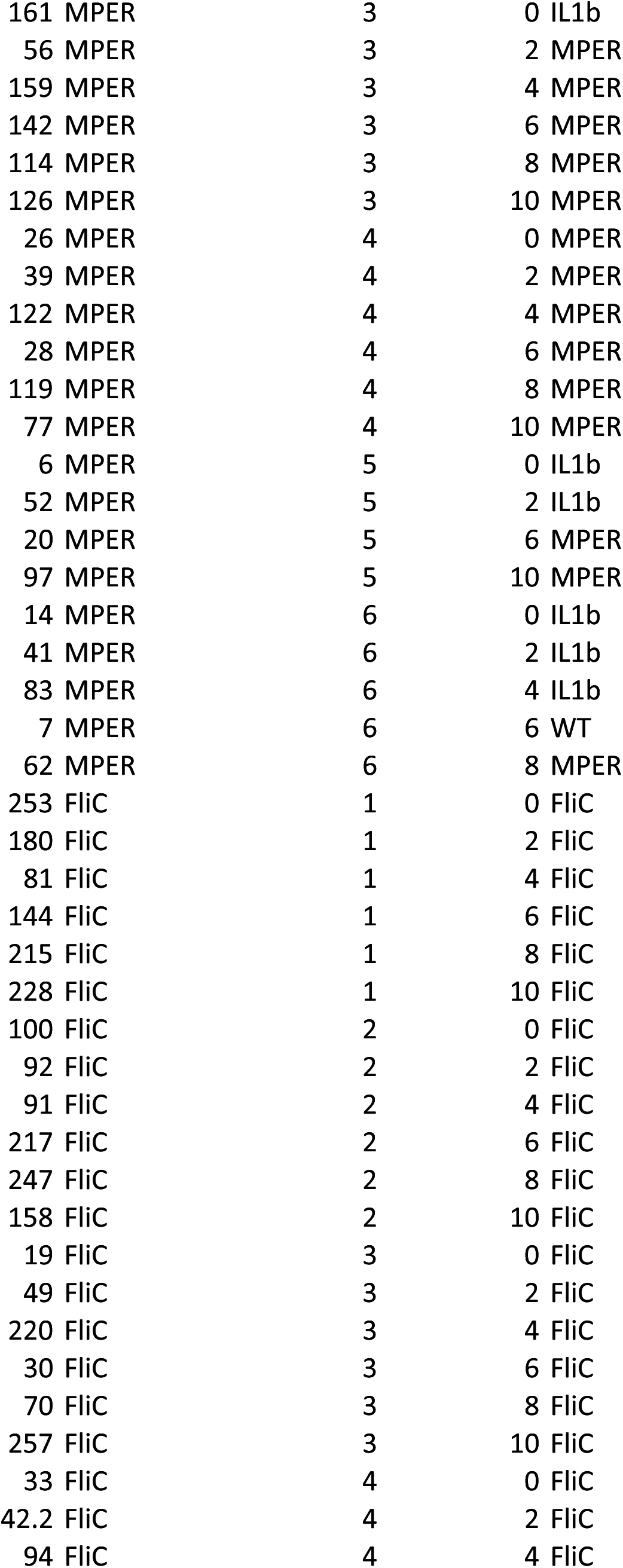
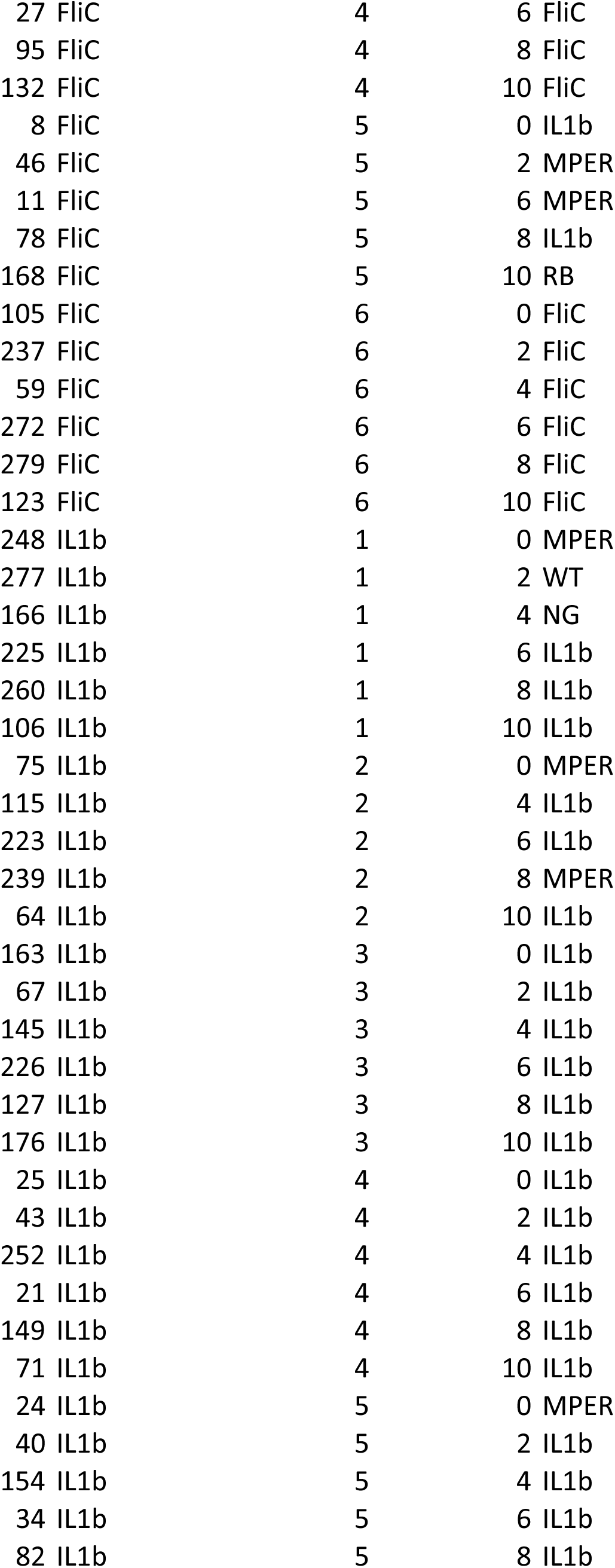
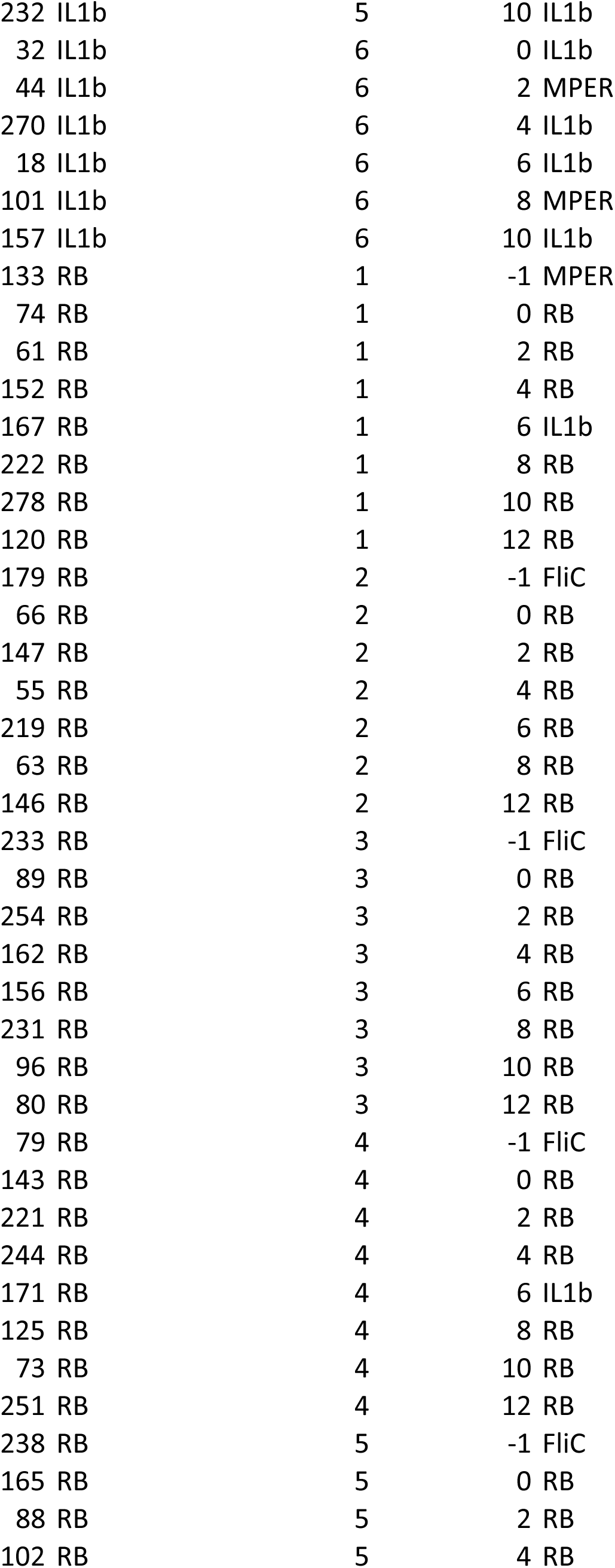
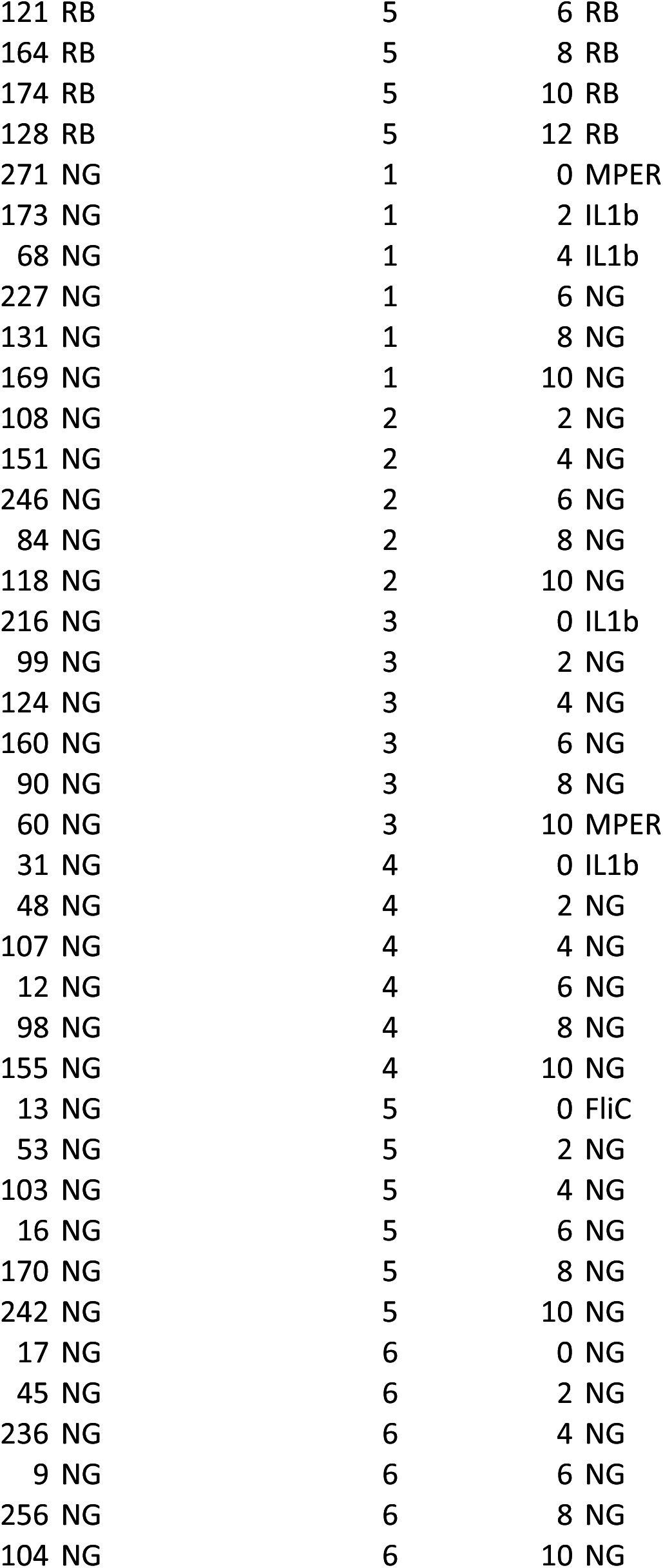
A comparison between observed and Random-Forest-predicted treatment classification per time and mouse (mouse.id). It is clear from this table that many misclassifications occur at time 0.

**Table S4:**
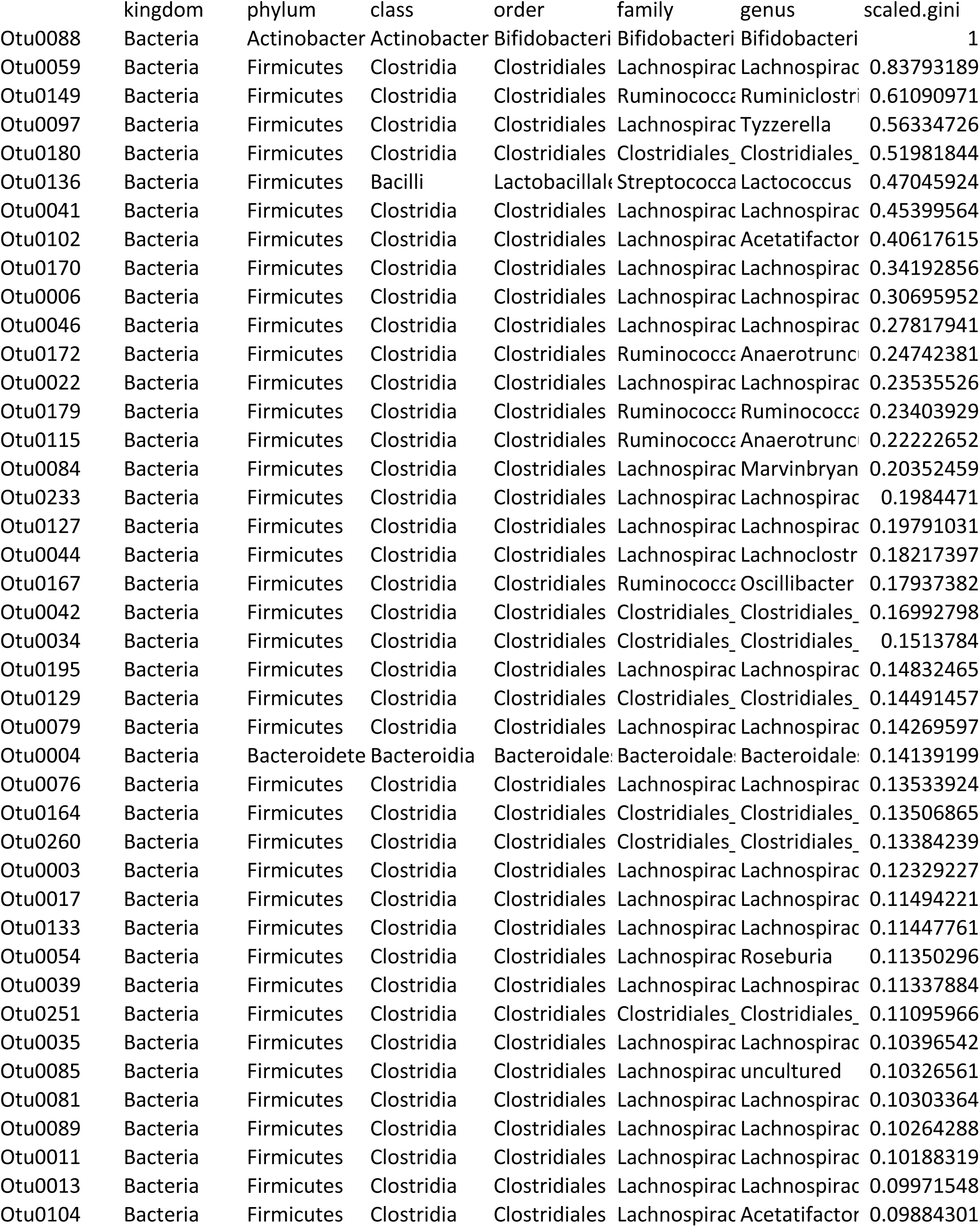
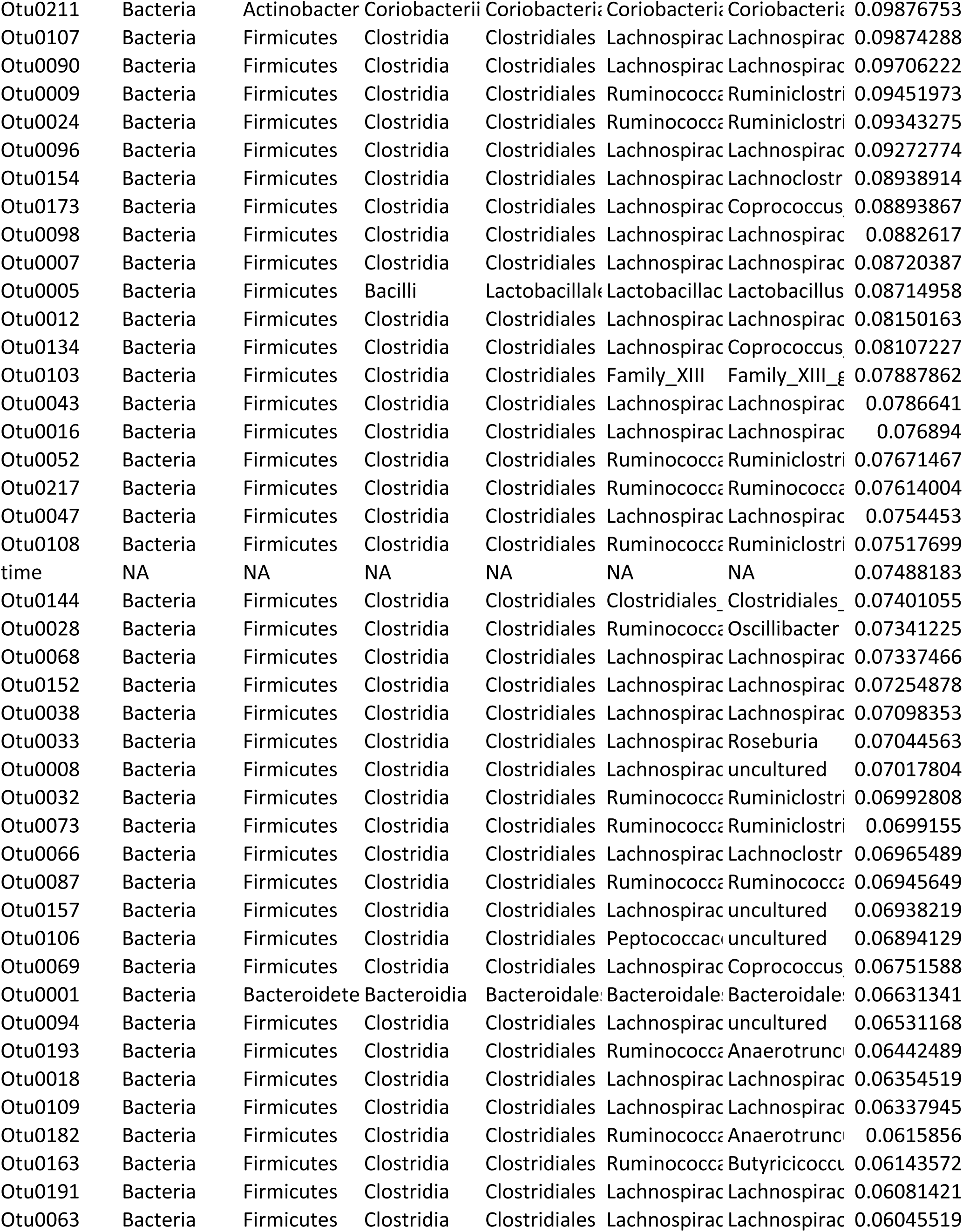
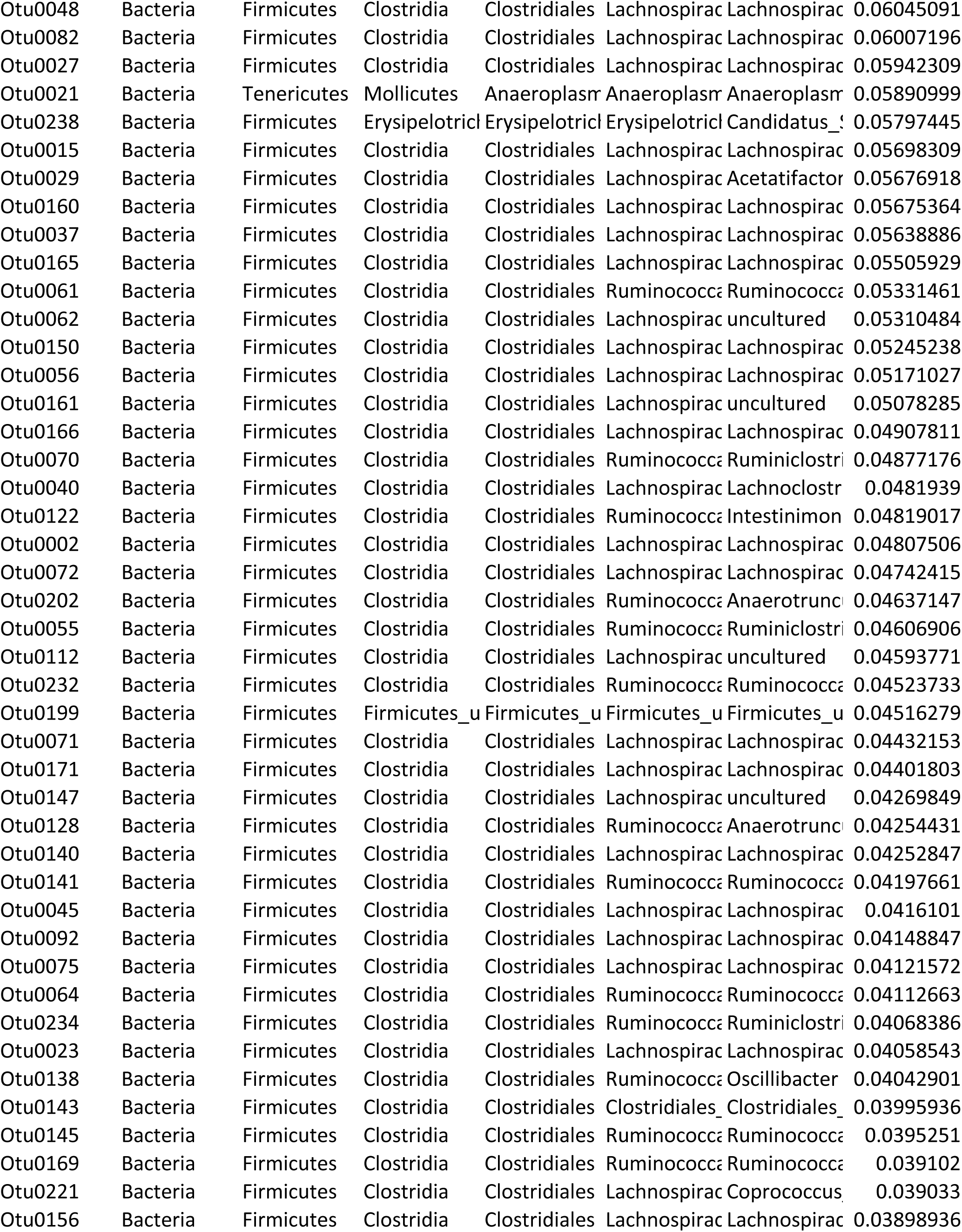
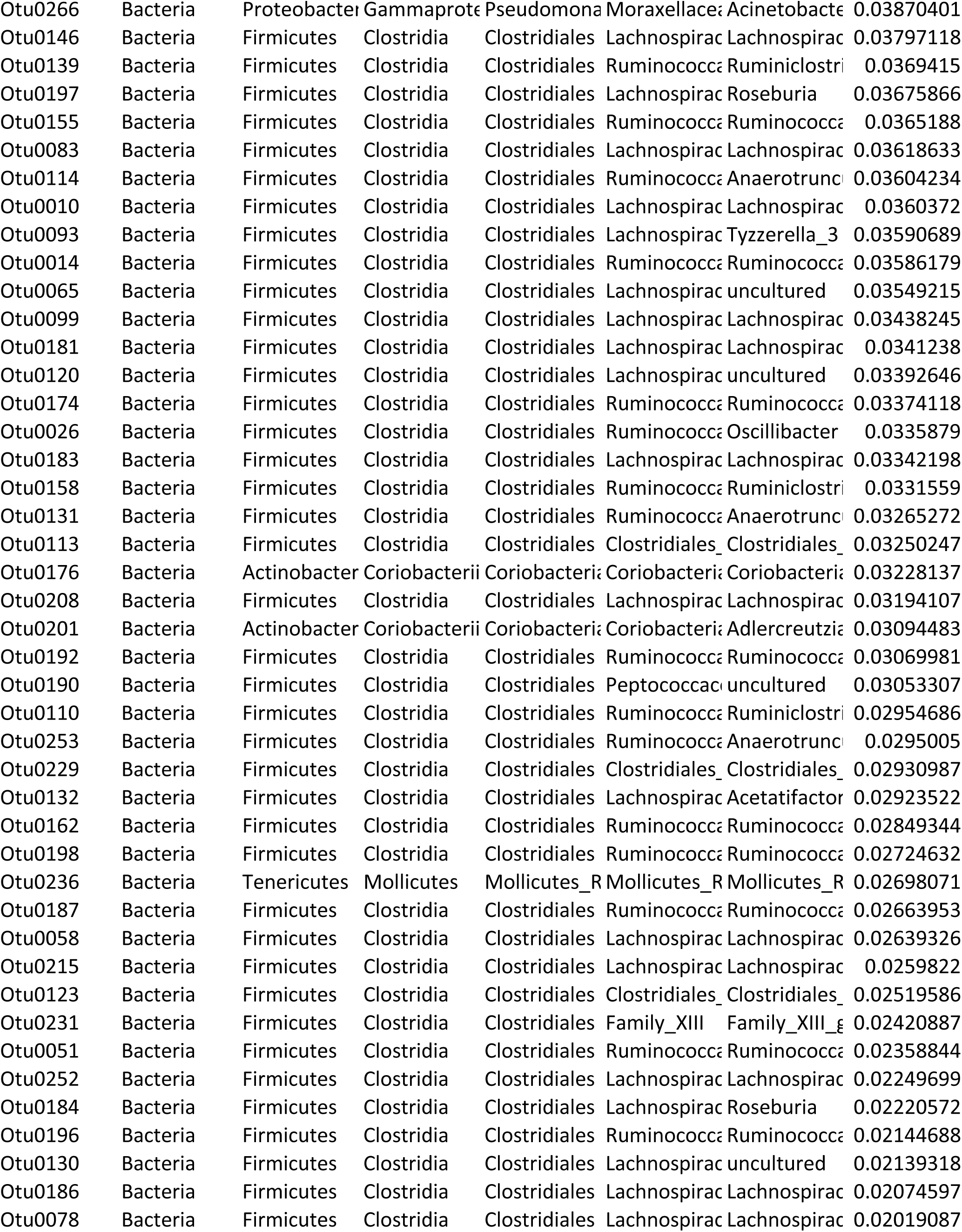
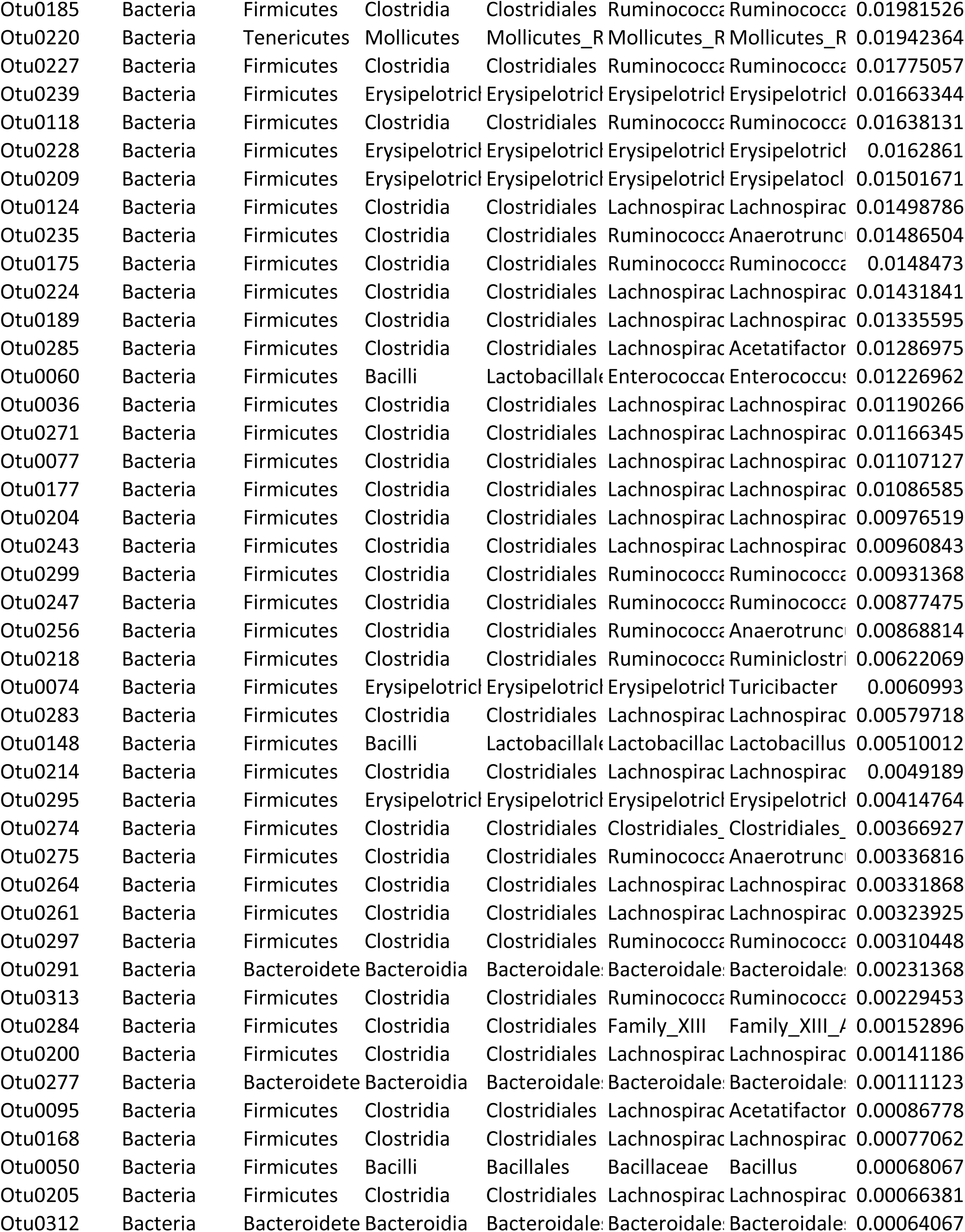
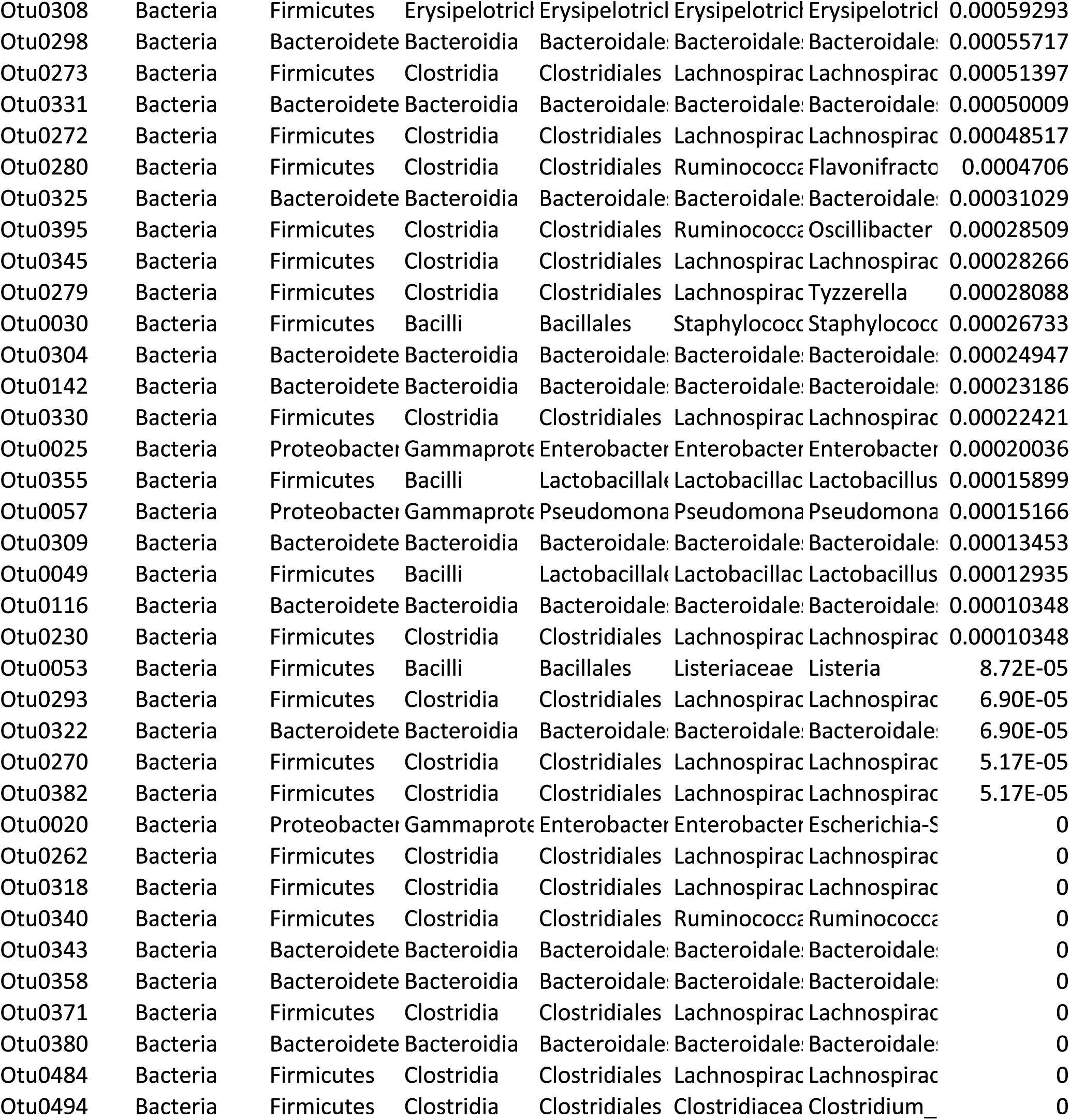
Per OTU-scaled GINI importance highlighting impact of each OTU (scaled.gini column) on the predicted classification of the fecal samples as obtained from the Random Forest. 1 is high and importance reduced as the number approaches 0.

**Table S5:** Confusion Matrix and out of bag (OOB) error rate of Random Forest classification of the cecal samples. Numbers on the diagonal of the table correspond to correctly classified samples while the off diagonal numbers represent misclassified samples. Rate of misclassification per class (treatment) is captured in the “class.error” column.

|  | FliC | IL1b | MPER | NG | RB | WT | class.error |
| --- | --- | --- | --- | --- | --- | --- | --- |
| FliC | 5 | 1 | 0 | 0 | 0 | 0 | 0.1666667 |
| IL1b | 0 | 5 | 1 | 0 | 0 | 0 | 0.1666667 |
| MPER | 0 | 1 | 3 | 1 | 0 | 1 | 0.5000000 |
| NG | 0 | 1 | 0 | 5 | 0 | 0 | 0.1666667 |
| RB | 0 | 0 | 0 | 0 | 4 | 0 | 0.0000000 |
| WT | 0 | 0 | 1 | 0 | 0 | 5 | 0.1666667 |

**Table S6:**
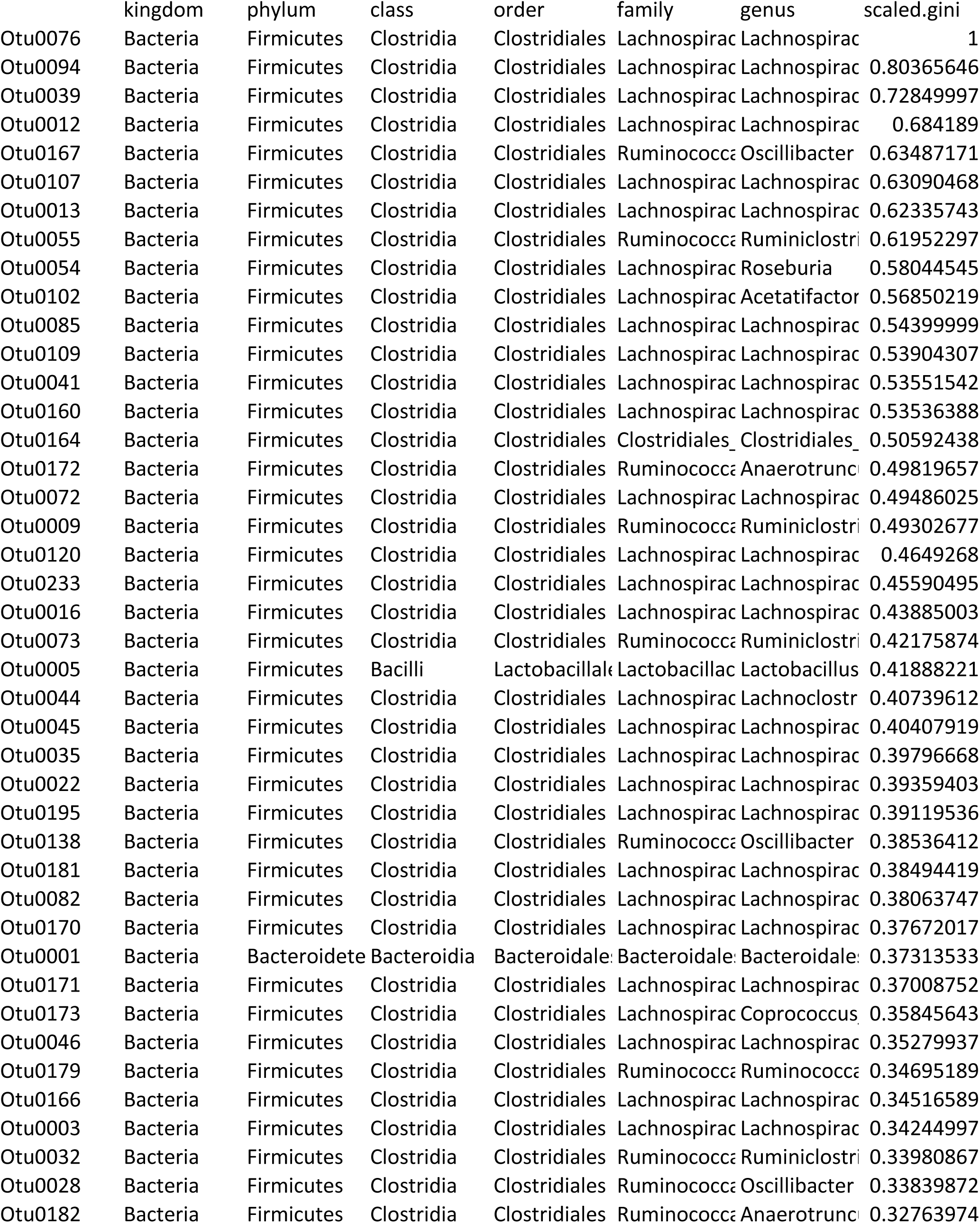
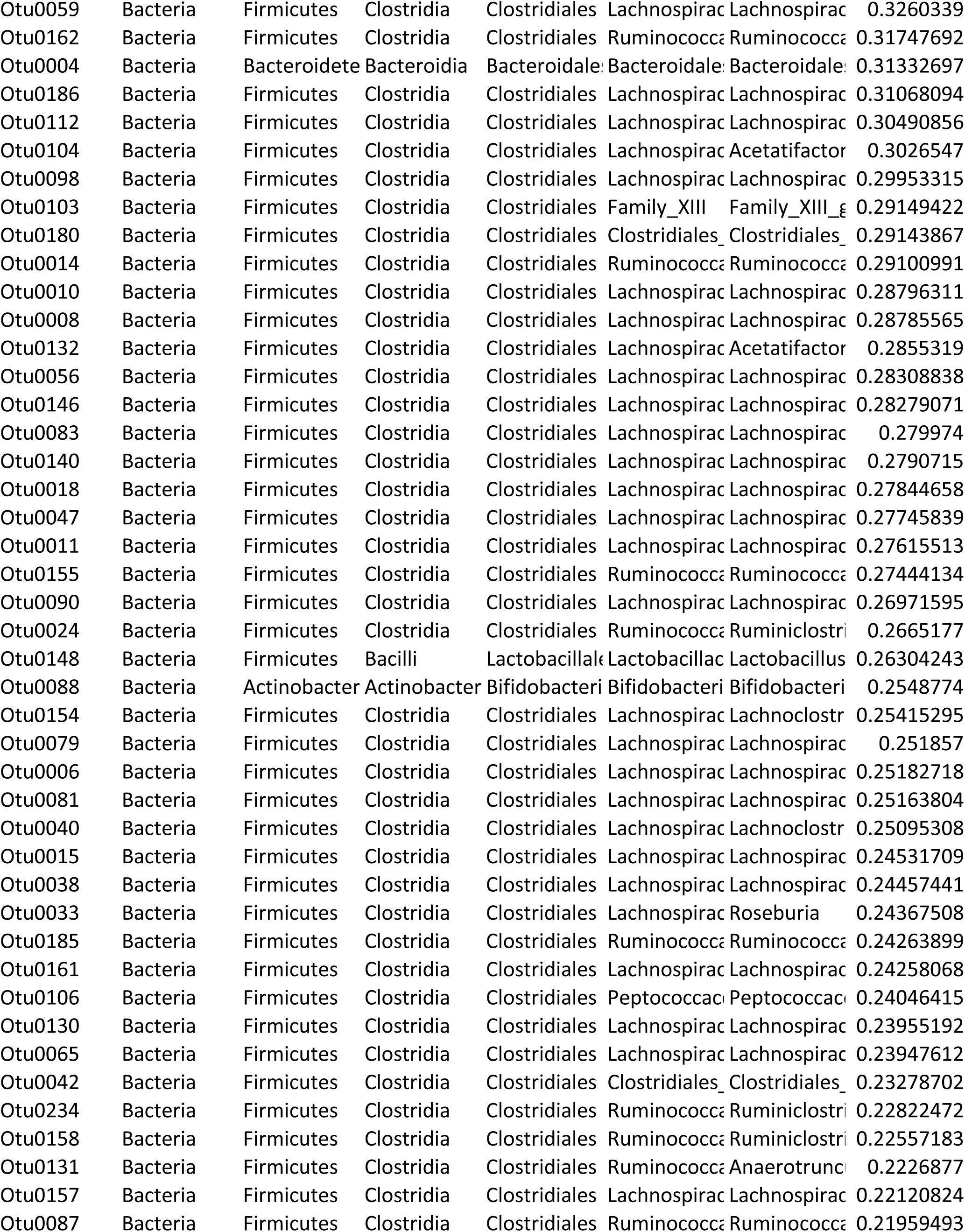
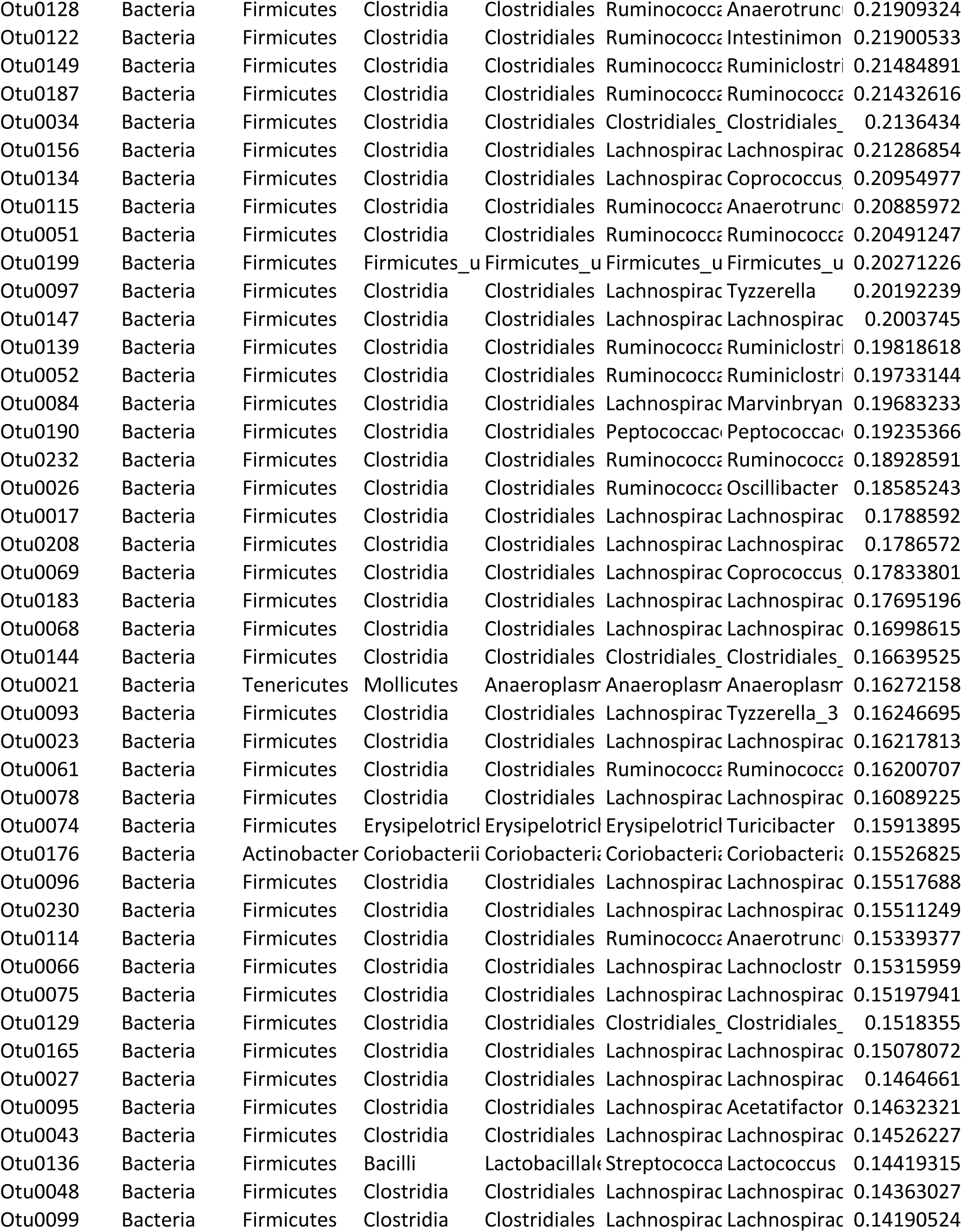
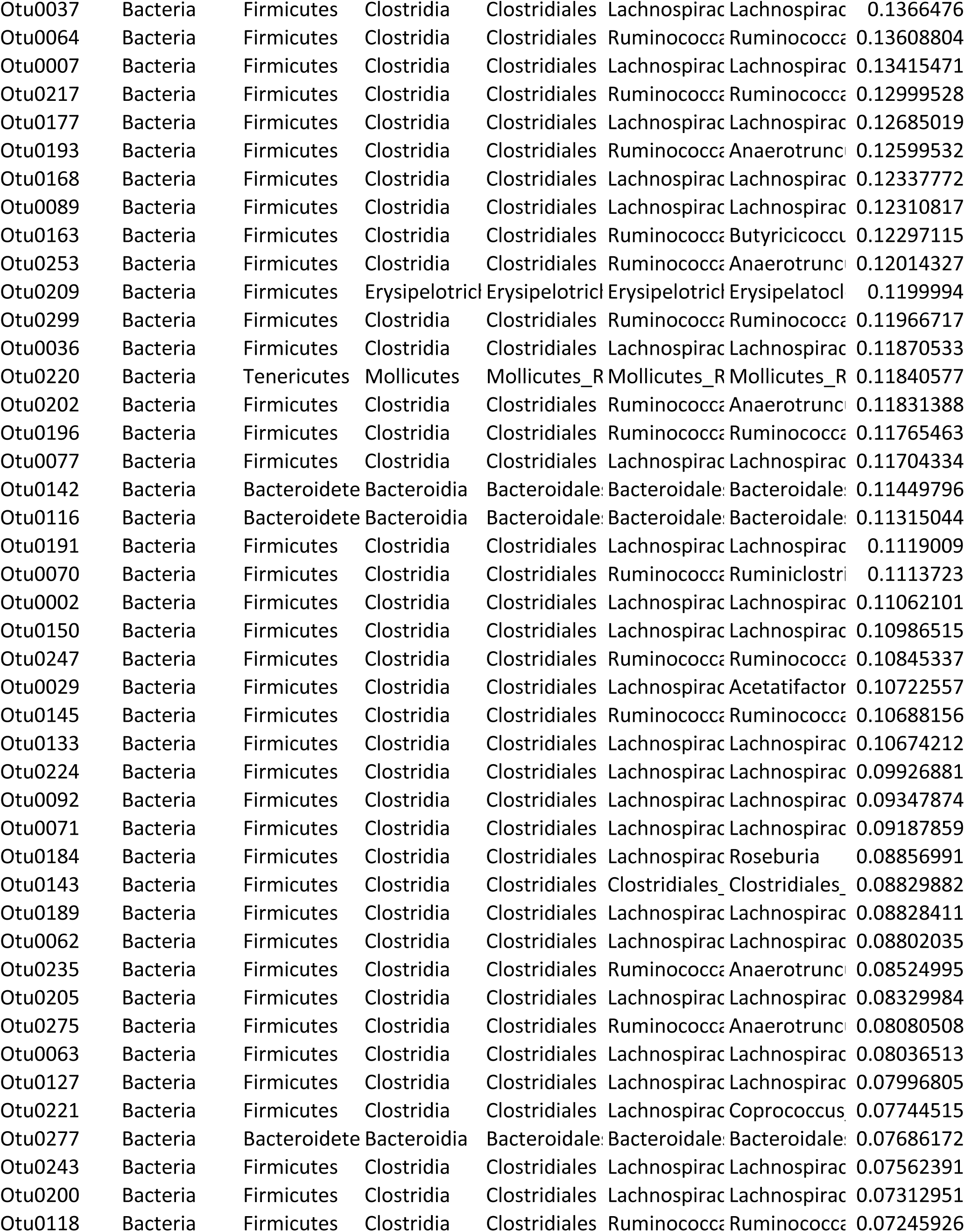
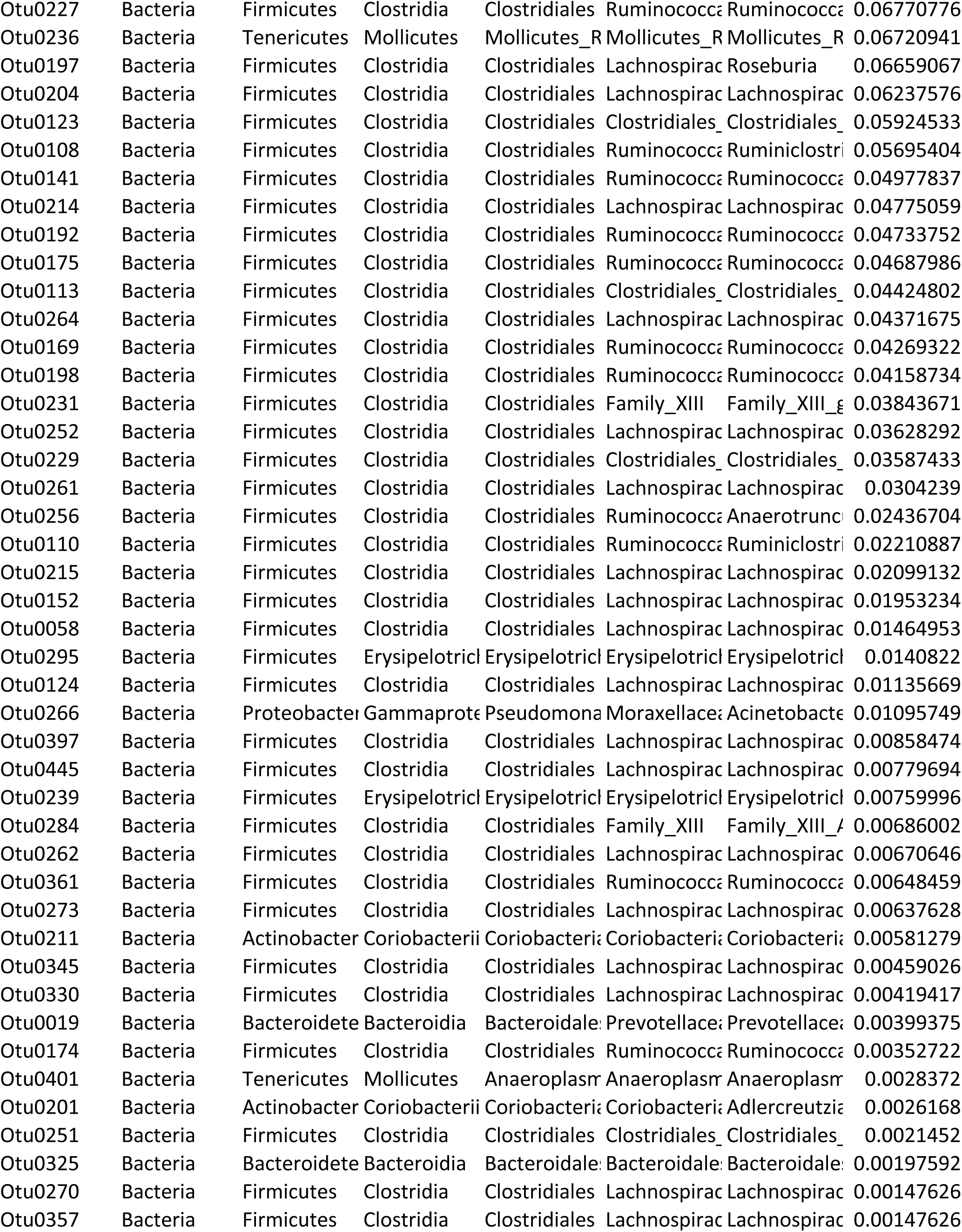
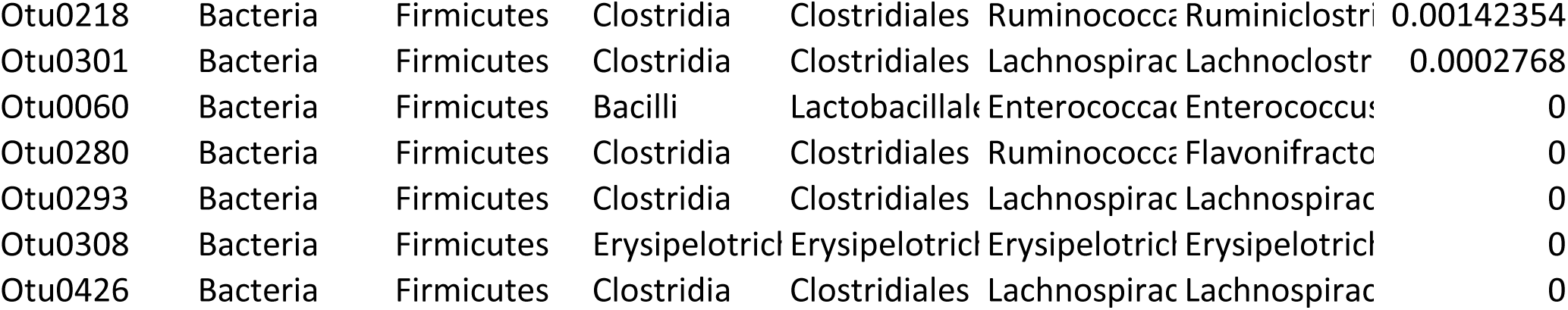
Per OTU-scaled GINI importance highlighting impact of each OTU (scaled.gini column) on the predicted classification of the cecal samples as obtained from the Random Forest. 1 is high and importance reduced as the number approaches 0.

**Table S7:**
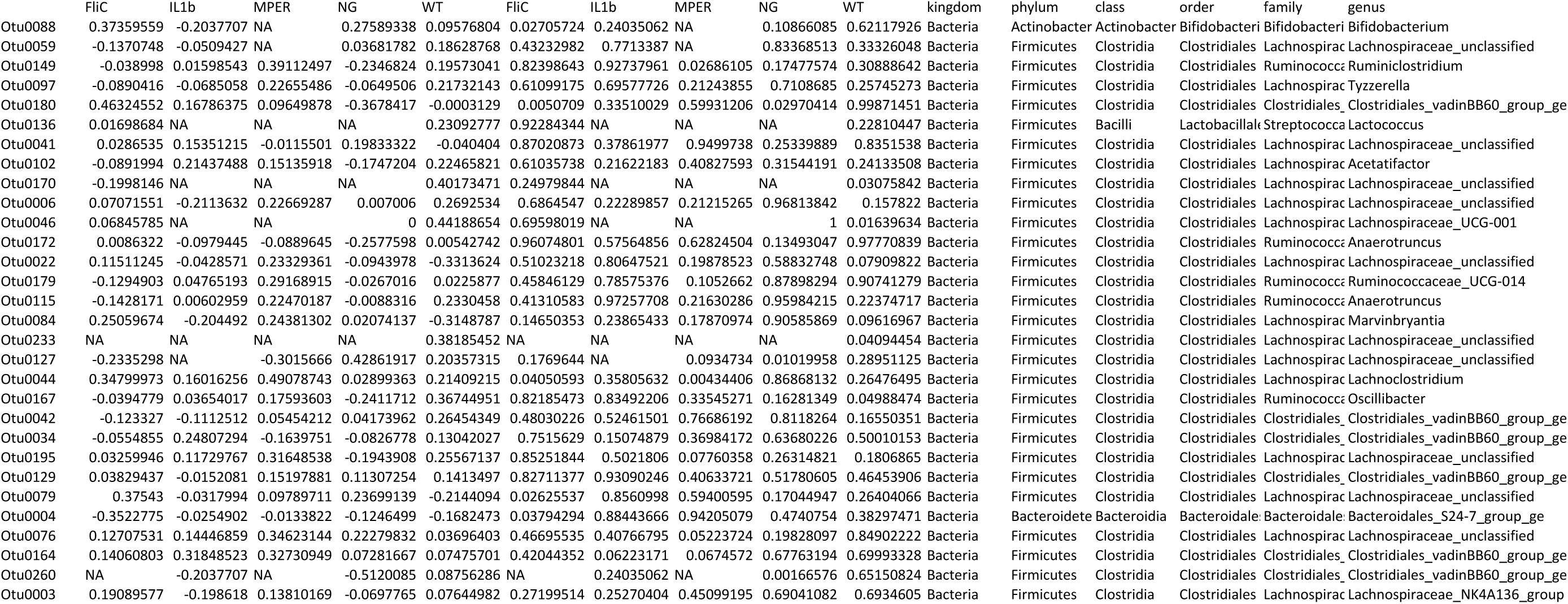
Spearman correlation between total IgA and abundance of the 30 most impactful OTUs based on the random forest analysis.

**Table S8:** Spearman correlation between MPER-specific IgA and abundance of the 30 most impactful OTUs based on the random forest analysis.

|  | FliC | IL1b | MPER | WT | FliC | IL1b | MPER | WT |
| --- | --- | --- | --- | --- | --- | --- | --- | --- |
| Otu0088 | NA | 0.11399367 | NA | -0.1112965 | NA | 0.55599771 | NA | 0.61315948 |
| Otu0059 | 0.22074238 | 0.11399367 | NA | -0.0161145 | 0.29995066 | 0.55599771 | NA | 0.9418245 |
| Otu0149 | -0.1535191 | -0.310527 | 0.38067434 | -0.1769955 | 0.4738675 | 0.101109 | 0.05504165 | 0.41914116 |
| Otu0097 | 0.0279527 | 0.08873947 | 0.19872099 | 0.08113857 | 0.89683998 | 0.64712407 | 0.33045391 | 0.71284758 |
| Otu0180 | 0.04614296 | -0.0081151 | -0.1573489 | -0.1250708 | 0.8304695 | 0.96667438 | 0.4426854 | 0.56962132 |
| Otu0136 | -0.1062688 | NA | NA | -0.0657272 | 0.62115083 | NA | NA | 0.76573361 |
| Otu0041 | 0.2196962 | 0.01312252 | 0.18945903 | 0.22504412 | 0.30230531 | 0.94613491 | 0.35393822 | 0.30188211 |
| Otu0102 | 0.01228193 | 0.24640844 | -0.1796617 | 0 | 0.95457802 | 0.19755187 | 0.3798239 | 1 |
| Otu0170 | -0.0416045 | NA | NA | 0.0656763 | 0.84694157 | NA | NA | 0.76591 |
| Otu0006 | 0.40028402 | 0.18486147 | 0.08464162 | 0.32838149 | 0.05259632 | 0.33704086 | 0.6809975 | 0.12606053 |
| Otu0046 | -0.3106903 | NA | NA | 0.09726833 | 0.13949713 | NA | NA | 0.65883808 |
| Otu0172 | 0.01228193 | -0.0683129 | -0.04355 | -0.0965669 | 0.95457802 | 0.72475873 | 0.83269931 | 0.66115511 |
| Otu0022 | -0.0343734 | 0.10911069 | 0.4376787 | 0 | 0.87331313 | 0.57314915 | 0.02534241 | 1 |
| Otu0179 | 0.31775996 | 0.03915396 | -0.0141917 | -0.0454545 | 0.13024695 | 0.84018812 | 0.94514231 | 0.8368365 |
| Otu0115 | 0.15628268 | 0.14269272 | 0.41372499 | 0.06437793 | 0.4658468 | 0.46026276 | 0.03563864 | 0.77041605 |
| Otu0084 | 0.24691286 | -0.1748747 | 0.2243268 | 0.29824736 | 0.24476195 | 0.36423536 | 0.27058932 | 0.16688656 |
| Otu0233 | NA | NA | NA | -0.2608287 | NA | NA | NA | 0.22933331 |
| Otu0127 | 0.03306673 | NA | -0.1198458 | -0.0657272 | 0.87809317 | NA | 0.55979226 | 0.76573361 |
| Otu0044 | 0.19330811 | 0.04340548 | 0.47112292 | 0.14467122 | 0.36544642 | 0.82309272 | 0.01512521 | 0.5101491 |
| Otu0167 | 0.01228193 | 0.00055376 | 0.36004375 | 0.17739127 | 0.95457802 | 0.99772532 | 0.07080553 | 0.41808493 |
| Otu0042 | 0.3223906 | -0.0081317 | -0.205046 | 0 | 0.12443311 | 0.96660626 | 0.31497287 | 1 |
| Otu0034 | 0.15522641 | -0.1327447 | -0.0961729 | 0.19380723 | 0.46890409 | 0.49242619 | 0.64024536 | 0.37557575 |
| Otu0195 | -0.1535191 | 0.06099629 | 0.23363413 | 0.03226098 | 0.4738675 | 0.75327868 | 0.25067956 | 0.88382058 |
| Otu0129 | -0.0698973 | -0.1136304 | 0.09247518 | -0.2184629 | 0.74552708 | 0.55726548 | 0.6532101 | 0.31661007 |
| Otu0079 | 0.18938343 | 0.09394121 | 0.29556138 | -0.3378168 | 0.37544704 | 0.62788173 | 0.14266575 | 0.11490604 |
| Otu0004 | 0.17305678 | -0.0508262 | 0.07331228 | -0.1125221 | 0.41870402 | 0.793447 | 0.72190398 | 0.60923092 |
| Otu0076 | 0.19387963 | 0.23803898 | 0.03821877 | -0.0964475 | 0.36400315 | 0.2136982 | 0.8529457 | 0.66154985 |
| Otu0164 | -0.1921209 | 0.04035438 | -0.1219464 | 0.30682766 | 0.36845503 | 0.83535369 | 0.55288143 | 0.15443113 |
| Otu0260 | NA | 0.11399367 | NA | -0.1113492 | NA | 0.55599771 | NA | 0.61299007 |
| Otu0003 | 0.1994247 | -0.0787474 | 0.49600781 | -0.1114021 | 0.35017266 | 0.68470906 | 0.00996532 | 0.61282044 |

## References

1. Sekirov, I., Russell, S. L., Antunes, L. C. M. & Finlay, B. B. Gut Microbiota in Health and Disease. Physiol. Rev. 90, 859–904 (2010).

2. Kinross, J. M., Darzi, A. W. & Nicholson, J. K. Gut microbiome-host interactions in health and disease. Genome Med 3, 14 (2011).

3. Cho, I. & Blaser, M. J. The human microbiome: at the interface of health and disease. Nat. Rev. Genet. 13, nrg3182 (2012).

4. Ley, R. E. et al. Obesity alters gut microbial ecology. Proc. Natl. Acad. Sci. 102, 11070–11075 (2005).

5. Bäckhed, F. et al. Defining a healthy human gut microbiome: current concepts, future directions, and clinical applications. Cell Host Microbe 12, (2012).

6. Bokulich, N. A. et al. Antibiotics, birth mode, and diet shape microbiome maturation during early life. Sci. Transl. Med. 8, 343ra82–343ra82 (2016).

7. Preidis, G. A. & Versalovic, J. Targeting the Human Microbiome With Antibiotics, Probiotics, and Prebiotics: Gastroenterology Enters the Metagenomics Era. Gastroenterology 136, 2015–2031 (2009).

8. Raymond, F. et al. The initial state of the human gut microbiome determines its reshaping by antibiotics. ISME J. (2015).

9. Petschow, B. et al. Probiotics, prebiotics, and the host microbiome: the science of translation: Probiotics, prebiotics, and the host microbiome. Ann. N. Y. Acad. Sci. 1306, 1–17 (2013).

10. Saulnier, D. M. et al. The intestinal microbiome, probiotics and prebiotics in neurogastroenterology. Gut Microbes 4, 17–27 (2013).

11. Sheflin, A. M. et al. Pilot Dietary Intervention with Heat-Stabilized Rice Bran Modulates Stool Microbiota and Metabolites in Healthy Adults. Nutrients 7, 1282–1300 (2015).

12. Gerritsen, J., Smidt, H., Rijkers, G. T. & Vos, W. M. Intestinal microbiota in human health and disease: the impact of probiotics. Genes Nutr. 6, 209 (2011).

13. Ng, S. C., Hart, A. L., Kamm, M. A., Stagg, A. J. & Knight, S. C. Mechanisms of Action of Probiotics: Recent Advances. Inflamm. Bowel Dis. 15, 300–310 (2009).

14. Sonnenburg, J. L., Chen, C. T. L. & Gordon, J. I. Genomic and Metabolic Studies of the Impact of Probiotics on a Model Gut Symbiont and Host. PLOS Biol. 4, e413 (2006).

15. O’Toole, P. W. & Cooney, J. C. Probiotic Bacteria Influence the Composition and Function of the Intestinal Microbiota. Interdiscip. Perspect. Infect. Dis. 9 (2008). doi:10.1155/2008/175285

16. Vanhoutvin, S. A. L. W. et al. Butyrate-Induced Transcriptional Changes in Human Colonic Mucosa. PLOS ONE 4, e6759 (2009).

17. Valdez, Y., Brown, E. M. & Finlay, B. B. Influence of the microbiota on vaccine effectiveness. Trends Immunol. 35, 526–537 (2014).

18. Harris, V. et al. Rotavirus vaccine response correlates with the infant gut microbiota composition in Pakistan. Gut Microbes 0, 1–9 (2017).

19. Harris, V. C. et al. Significant Correlation Between the Infant Gut Microbiome and Rotavirus Vaccine Response in Rural Ghana. J. Infect. Dis. 215, 34–41 (2017).

20. Kandasamy, S., Chattha, K. S., Vlasova, A. N., Rajashekara, G. & Saif, L. J. Lactobacilli and Bifidobacteria enhance mucosal B cell responses and differentially modulate systemic antibody responses to an oral human rotavirus vaccine in a neonatal gnotobiotic pig disease model. Gut Microbes 5, 639–651 (2014).

21. Marelli, B., Perez, A. R., Banchio, C., de Mendoza, D. & Magni, C. Oral immunization with live Lactococcus lactis expressing rotavirus VP8* subunit induces specific immune response in mice. J. Virol. Methods 175, 28–37 (2011).

22. Pant, N., Marcotte, H., Brüssow, H., Svensson, L. & Hammarström, L. Effective prophylaxis against rotavirus diarrhea using a combination of Lactobacillus rhamnosus GG and antibodies. BMC Microbiol. 7, 86 (2007).

23. Park, M. S., Kwon, B., Ku, S. & Ji, G. E. The Efficacy of Bifidobacterium longum BORI and Lactobacillus acidophilus AD031 Probiotic Treatment in Infants with Rotavirus Infection. Nutrients 9, 887 (2017).

24. Lee, I.-C., Tomita, S., Kleerebezem, M. & Bron, P. A. The quest for probiotic effector molecules—Unraveling strain specificity at the molecular level. Pharmacol. Res. 69, 61–74 (2013).

25. Tarahomjoo, S. Development of Vaccine Delivery Vehicles Based on Lactic Acid Bacteria. Mol. Biotechnol. 51, 183–199 (2012).

26. LeBlanc, J. G. et al. Mucosal targeting of therapeutic molecules using genetically modified lactic acid bacteria: an update. FEMS Microbiol. Lett. 344, 1–9 (2013).

27. LeCureux, J. S. & Dean, G. A. Lactobacillus Mucosal Vaccine Vectors: Immune Responses against Bacterial and Viral Antigens. mSphere 3, e00061–18 (2018).

28. Ruiz, L., Margolles, A. & Sánchez, B. Bile resistance mechanisms in Lactobacillus and Bifidobacterium. Front. Microbiol. 4, (2013).

29. Kajikawa, A. et al. Dissimilar Properties of Two Recombinant Lactobacillus acidophilus Strains Displaying Salmonella FliC with Different Anchoring Motifs. Appl. Environ. Microbiol. 77, 6587–6596 (2011).

30. Wells, J. M. & Mercenier, A. Mucosal delivery of therapeutic and prophylactic molecules using lactic acid bacteria. Nat. Rev. Microbiol. 6, 349–362 (2008).

31. Lebeer, S., Vanderleyden, J. & De Keersmaecker, S. C. J. Host interactions of probiotic bacterial surface molecules: comparison with commensals and pathogens. Nat. Rev. Microbiol. 8, 171–184 (2010).

32. Girardin, S. E. et al. Nod2 Is a General Sensor of Peptidoglycan through Muramyl Dipeptide (MDP) Detection. J. Biol. Chem. 278, 8869–8872 (2003).

33. Matsuguchi, T. et al. Lipoteichoic Acids from Lactobacillus Strains Elicit Strong Tumor Necrosis Factor Alpha-Inducing Activities in Macrophages through Toll-Like Receptor 2. Clin Diagn Lab Immunol 10, 259–266 (2003).

34. Zeuthen, L. H., Fink, L. N. & Frøkiær, H. Toll-like receptor 2 and nucleotide-binding oligomerization domain-2 play divergent roles in the recognition of gut-derived lactobacilli and bifidobacteria in dendritic cells. Immunology 124, 489–502 (2008).

35. Chorny, A., Puga, I. & Cerutti, A. Chapter 2 - Innate Signaling Networks in Mucosal IgA Class Switching. in Advances in Immunology (eds. Fagarasan, S. & Cerutti, A.) 107, 31–69 (Academic Press, 2010).

36. Konstantinov, S. R. et al. S layer protein A of Lactobacillus acidophilus NCFM regulates immature dendritic cell and T cell functions. Proc. Natl. Acad. Sci. 105, 19474–19479 (2008).

37. Van Tassell, M. L. & Miller, M. J. Lactobacillus Adhesion to Mucus. Nutrients 3, 613–636 (2011).

38. Vélez, M. P., De Keersmaecker, S. C. J. & Vanderleyden, J. Adherence factors of Lactobacillus in the human gastrointestinal tract. FEMS Microbiol. Lett. 276, 140–148 (2007).

39. Kajikawa, A. et al. Mucosal Immunogenicity of Genetically Modified Lactobacillus acidophilus Expressing an HIV-1 Epitope within the Surface Layer Protein. PLOS ONE 10, e0141713 (2015).

40. Kajikawa, A., Masuda, K., Katoh, M. & Igimi, S. Adjuvant Effects for Oral Immunization Provided by Recombinant Lactobacillus casei Secreting Biologically Active Murine Interleukin-1β. Clin Vaccine Immunol 17, 43–48 (2010).

41. Kajikawa, A. et al. Construction and Immunological Evaluation of Dual Cell Surface Display of HIV-1 Gag and Salmonella enterica Serovar Typhimurium FliC in Lactobacillus acidophilus for Vaccine Delivery. Clin Vaccine Immunol 19, 1374–1381 (2012).

42. Yang, X. et al. High protective efficacy of rice bran against human rotavirus diarrhea via enhancing probiotic growth, gut barrier function, and innate immunity. Sci. Rep. 5, 15004 (2015).

43. Chassaing, B., Ley, R. E. & Gewirtz, A. T. Intestinal Epithelial Cell Toll-like Receptor 5 Regulates the Intestinal Microbiota to Prevent Low-Grade Inflammation and Metabolic Syndrome in Mice. Gastroenterology 147, 1363–1377.e17 (2014).

44. Kajikawa, A. & Igimi, S. Development of Recombinant Vaccines in Lactobacilli for Elimination of Salmonella. Biosci. Microflora 30, 93–98 (2011).

45. Gerritsen, J., Smidt, H., Rijkers, G. T. & Vos, W. M. de. Intestinal microbiota in human health and disease: the impact of probiotics. Genes Nutr. 6, 209 (2011).

46. Cleland, E. J. et al. Probiotic manipulation of the chronic rhinosinusitis microbiome. Int. Forum Allergy Rhinol. 4, 309–314 (2014).

47. Kandasamy, S. et al. Unraveling the Differences between Gram-Positive and Gram-Negative Probiotics in Modulating Protective Immunity to Enteric Infections. Front. Immunol. 8, (2017).

48. Harris, V. C. The Significance of the Intestinal Microbiome for Vaccinology: From Correlations to Therapeutic Applications. Drugs 78, 1063–1072 (2018).

49. Parker, E. P. K. et al. Influence of the intestinal microbiota on the immunogenicity of oral rotavirus vaccine given to infants in south India. Vaccine 36, 264–272 (2018).

50. Parker, E. P. et al. Causes of impaired oral vaccine efficacy in developing countries. Future Microbiol. 13, 97–118 (2018).

51. Sheflin, A. M. et al. Dietary supplementation with rice bran or navy bean alters gut bacterial metabolism in colorectal cancer survivors. Mol. Nutr. Food Res. 61, 1500905 (2017).

52. Broussard, J. L. & Devkota, S. The changing microbial landscape of Western society: Diet, dwellings and discordance. Mol. Metab. 5, 737–742 (2016).

53. Smits, S. A., Marcobal, A., Higginbottom, S., Sonnenburg, J. L. & Kashyap, P. C. Individualized Responses of Gut Microbiota to Dietary Intervention Modeled in Humanized Mice. mSystems 1, e00098–16 (2016).

54. Bisanz, J. E., Upadhyay, V., Turnbaugh, J. A., Ly, K. & Turnbaugh, P. Diet Induces Reproducible Alterations in the Mouse and Human Gut Microbiome. (Social Science Research Network, 2019).

55. Meehan, C. J. & Beiko, R. G. A Phylogenomic View of Ecological Specialization in the Lachnospiraceae, a Family of Digestive Tract-Associated Bacteria. Genome Biol. Evol. 6, 703–713 (2014).

56. Milani, C. et al. Bifidobacteria exhibit social behavior through carbohydrate resource sharing in the gut. Sci. Rep. 5, 15782 (2015).

57. Wang, X., Brown, I. L., Evans, A. J. & Conway, P. L. The protective effects of high amylose maize (amylomaize) starch granules on the survival of Bifidobacterium spp. in the mouse intestinal tract. J. Appl. Microbiol. 87, 631–639 (1999).

58. Laukens, D., Brinkman, B. M., Raes, J., De Vos, M. & Vandenabeele, P. Heterogeneity of the gut microbiome in mice: guidelines for optimizing experimental design. FEMS Microbiol. Rev. 40, 117–132 (2016).

59. Kumar, A. et al. Dietary rice bran promotes resistance to Salmonella enterica serovar Typhimurium colonization in mice. BMC Microbiol. 12, 71 (2012).

60. Kajikawa, A. et al. Mucosal Immunogenicity of Genetically Modified Lactobacillus acidophilus Expressing an HIV-1 Epitope within the Surface Layer Protein. PLOS ONE 10, e0141713 (2015).

61. Bumgardner, S. A. et al. Nod2 is required for antigen-specific humoral responses against antigens orally delivered using a recombinant Lactobacillus vaccine platform. PLOS ONE 13, e0196950 (2018).

62. Stoeker, L. et al. Assessment of Lactobacillus gasseri as a Candidate Oral Vaccine Vector. Clin. Vaccine Immunol. 18, 1834–1844 (2011).

63. Frey, A., Di Canzio, J. & Zurakowski, D. A statistically defined endpoint titer determination method for immunoassays. J. Immunol. Methods 221, 35–41 (1998).

64. Methé, B. A. et al. A framework for human microbiome research. Nature 486, 215–221 (2012).

65. Kozich, J. J., Westcott, S. L., Baxter, N. T., Highlander, S. K. & Schloss, P. D. Development of a Dual-Index Sequencing Strategy and Curation Pipeline for Analyzing Amplicon Sequence Data on the MiSeq Illumina Sequencing Platform. Appl. Environ. Microbiol. 79, 5112–5120 (2013).

66. Schloss, P. D., et al. Introducing mothur: Open-Source, Platform-Independent, Community-Supported Software for Describing and Comparing Microbial Communities. Appl. Environ. Microbiol. 75, 7537–7541 (2009).

67. Quast, C. et al. The SILVA ribosomal RNA gene database project: improved data processing and web-based tools. Nucleic Acids Res. 41, D590–D596 (2013).

68. Westcott, S. L. & Schloss, P. D. OptiClust, an Improved Method for Assigning Amplicon-Based Sequence Data to Operational Taxonomic Units. mSphere 2, e00073–17 (2017).

69. Oksanen, J., et al. vegan: *Community Ecology Package*. (2014).

70. R Core Team. R: A language and environment for statistical computing. (2017).

71. McMurdie, P. J. & Holmes, S. phyloseq: An R Package for Reproducible Interactive Analysis and Graphics of Microbiome Census Data. PLOS ONE 8, e61217 (2013).

72. Plummer, M., Stukalov, A. & Denwood, M. Package ‘rjags’. (2016).

73. Plummer, M. & others. JAGS: A program for analysis of Bayesian graphical models using Gibbs sampling. in Proceedings of the 3rd international workshop on distributed statistical computing 124, 125 (Vienna, 2003).

74. Spiegelhalter, D. J., Best, N. G., Carlin, B. P. & Van Der Linde, A. Bayesian measures of model complexity and fit. J. R. Stat. Soc. Ser. B Stat. Methodol. 64, 583–639 (2002).

75. Gelman, A. & Rubin, D. B. Inference from iterative simulation using multiple sequences. Stat. Sci. 7, 457–511 (1992).

76. Legendre, P. & Legendre, L. Numerical Ecology. Vol. 20, (Elsevier B.V., 1998).

77. Paulson, J. N., Stine, O. C., Bravo, H. C. & Pop, M. Differential abundance analysis for microbial marker-gene surveys. Nat. Methods 10, 1200–1202 (2013).

78. Wickham, H. ggplot2: Elegant graphics for data analysis. (Springer Science+Business Media, 2009).

79. Breiman, L. Random forests. Mach. Learn. 45, 5–32 (2001).

80. Breiman, L., Cutler, A., Liaw, A. & Wiener, M. Package ‘randomForest’. (2015).

81. Spearman, C. Correlation between arrays in a table of correlations. Proc. R. Soc. Lond. Ser. Contain. Pap. Math. Phys. Character 101, 94–100 (1922).

